# Insects fight low-dose infections with terminal investment, not innate immunity

**DOI:** 10.1101/2025.04.03.646978

**Authors:** Jingbo Liu, Nicholas K. Priest

## Abstract

Though the innate immune system is considered to be the primary defence promoting survival against pathogenic microbes, non-immunological strategies may provide cost-effective responses against ubiquitous low-dose infections. However, research using model systems has yet to develop a framework for systematically varying topical exposure doses, much less for examining how hosts mitigate the fitness costs associated with immune deployment. Here, we demonstrate that insects respond to low-dose infections with terminal investment. Female fruit flies, *Drosophila melanogaster*, have higher egg-to adult viability and lower survival when exposed to low-dose or sexually transmitted infections of an endemic fungus, *Aspergillus austwickii*. We identify *Turandot* C (*TotC*), a humoral stress response gene in insects, as the first non-immunological regulator of fecundity compensation. Strikingly, *TotC*-mediated fecundity compensation imposes negligible lifetime fitness costs, whereas expression of the canonical immune gene *Dorsal-related immunity factor* (*Dif*) triggers reproductively costly antagonistic pleiotropy. Contrary to foundational ecological theory, we show that terminal investment arises from immediate survival-reproduction trade-offs, not truncated reproductive potential. Our findings reveal that adaptive evolution of innate immunity is constrained by classical fitness trade-offs in response to ubiquitous low-dose infections, which permits the evolution of mechanisms in which the host strategically surrenders to the pathogen. Our study refines how we conceptualise host-pathogen evolutionary conflicts and underscores a need to understand how immunosuppression evolves in hosts under ecologically prevalent low-dose infections.

## Introduction

Terminal investment theory posits the counterintuitive prediction that organisms respond to infections and other survival threats not by enhancing survival but by increasing reproductive success at the cost of survival or future reproductive output (Williams, 1966; Hirshfield and Tinkle, 1975; Pianka and Park, 1975; Duffield et al., 2017). Terminal investment induced by parasitic challenge has been extensively documented across the animal kingdom (Minchella and Loverde, 1981; Polak and Starmer, 1998; Bonneaud et al., 2004; Weil et al., 2006; Brannelly et al., 2016; Hudson et al., 2020; Schulz et al., 2023; Liao et al., 2024). These phenomena have been interpreted variously as surrender mechanisms, ‘mafia-like’ pathogen manipulations or adaptive non-immunological defences (Thomas et al., 2005; Parker et al., 2011). However, it remains unclear whether terminal investment is specific to low-dose infections and how it relates to innate immunity. Existing immunological research has predominantly focused on survival-enhancing mechanisms that promote resistance to lethal infections (Hoffmann and Reichhart, 2002; Moita et al., 2005; Hanson et al., 2023). In contrast, the evolutionary consequences of exposure to low-dose and sexually transmitted infections are not poorly understood, despite these representing some of the most common types of insect-pathogen interactions in nature (Knell and Webberley, 2004; Zhong et al. 2013; Adamo, 2014).

The dynamic threshold model of terminal investment proposes that infection load determines whether infected organisms adopt a high-survival, low-reproduction strategy or a low-survival, high-reproduction strategy, as resource limitations constrain optimal adjustments to both fitness traits simultaneously (Duffield et al., 2017). The evolutionary rationale behind terminal investment is that increased reproductive effort arises from ecological perturbations that reduce Residual Reproductive Value (RRV), a formal estimator of reproductive potential (Williams, 1966; Hirshfield and Tinkle, 1975; Pianka and Park, 1975; Clutton-Brock, 1984; Stearns, 1992). However, RRV is rarely estimated, and no study has tested the alternate hypothesis that terminal investment is driven by trade-offs acting synchronously (Duffield et al., 2017; Farchmin et al., 2020; Corbel and Carazo, 2022; Jehan et al., 2022; Saraux and Chiaradia, 2022; Schulz et al., 2023; Foo et al., 2023; but see Kolora et al., 2021). Ecological theory suggests that defence against infection is imperfect due to fitness trade-offs that prevent the optimal expression of defence traits, leading to negative correlations between fitness traits (Viney et al., 2005; McKean and Lazzaro, 2011). Although challenging to identify (see Van Noordwijk and De Jong, 1986; Reznick et al., 2000), evidence for negative genetic correlations between fitness traits under different infection loads could provide insights into the ecological contexts that underlie the adaptive value of terminal investment and innate immunity (Williams, 1957 & 1966; Kirkwood and Rose, 1991; Stearns, 1992; Brommer, 2000; Laskowski et al., 2021).

Insects may mitigate the risks of disease exposure through various mechanisms that manage different infection loads. One way to investigate this is by identifying candidate immunological and non-immunological genes and assessing how fitness trade-offs differ under varying topical infection loads. *Dorsal-related immunity factor* (*Dif*) is an ideal model for examining the fitness consequences of innate immunity genes, given its role in the Toll signalling pathway and its regulation of the costly production of antimicrobial peptides, which provide effective defence against topical fungal infections (Ip et al., 1993; Lemaitre et al., 1996). Similarly, *Turandot C* (*TotC*) represents a promising candidate for terminal investment in response to low-dose sexually transmitted infections (STIs). *Tot* genes are a rapidly evolving family of humoral stress-response factors in insects, whose function in managing environmental and pathogenic challenges is a common feature of large gene families broadly present in diverse species (Ekengren and Hultmark, 2001). Despite its high expression in response to mating calls from conspecific males, *TotC* does not enhance survival in females exposed to topical fungal infections, which implicates a potential role in reproduction (Immonen and Ritchie, 2012; Zhong et al., 2013). Recent research shows that gene products of *Turandot* genes reduce the cytotoxicity of the antimicrobial peptides, indicating that the expression of *TotC* and *Dif* have very different consequences for post-infection fitness (Rommelaere et al. 2024).

Here, we test the hypothesis that terminal investment evolved as a low-cost response to low-dose infections. We developed an innovative needle-tap method to topically inoculate insects with entomopathogenic fungi. To assess dose-specific variation in fitness trade-offs, we isolated a fungus, *Aspergillus austwickii*, from a lab-adapted strain, the Dahomey line of *Drosophila melanogaster*, and evaluated lifetime reproductive output and survival patterns in Dahomey under a gradient of topical infection doses of (including zero, one, three, sexually transmitted and direct topical inoculations). To test the prediction that *TotC* expression mediates reproductive responses to low-dose infections without substantially compromising lifetime reproductive success, we used the Gal4/UAS gene knockdown system and assessed lifetime fitness outcomes under zero, sexually transmitted, and direct topical inoculations of a model entomopathogenic fungus *Metarhizium robertsii* (chosen due to its identifiable role in causing death, unlike endemic fungi in fruit flies). Additionally, we tested whether *TotC* operates through stress-responsive pathways distinct from innate immunity by comparing the fitness consequences of *Dif* expression under the same infection conditions. In contrast to *TotC*, we predicted that *Dif* expression would be essential for protection against high-dose infections but ineffective, showing strong antagonistic pleiotropy between survival and reproduction, in response to low-dose sexually transmitted infections.

## Results

### Low-dose fungal infections induce terminal investment

We developed novel inoculation methods in *Drosophila melanogaster* using an indigenous fungus, *A. austwickii*, to assess the impact of low-dose infection on fitness (Figure 1). The microbial loads varied among the different infection modes, ranging from 6.29 ± 0.49 × 10³ ml⁻¹ for the One Dose treatment to 255 ± 6 × 10³ ml⁻¹ for the Direct Topical Infections (DTIs), compared to untreated naïve females at 0.933 × 10³ ± 0.166 ml⁻¹. Inoculation levels for the Three Doses and STI treatments were comparable, at 23.2 ± 1.6 × 10³ ml⁻¹ and 35.2 ± 3.0 × 10³ ml⁻¹, respectively. Physical evidence of topical inoculation is discernible under light microscopy for the needle-tap (Figure 1B, 1C, 1D) and DTI treatments (Figure 1F), though it is less apparent in females infected via sexual transmission (Figure 1E).

**Figure 1.**
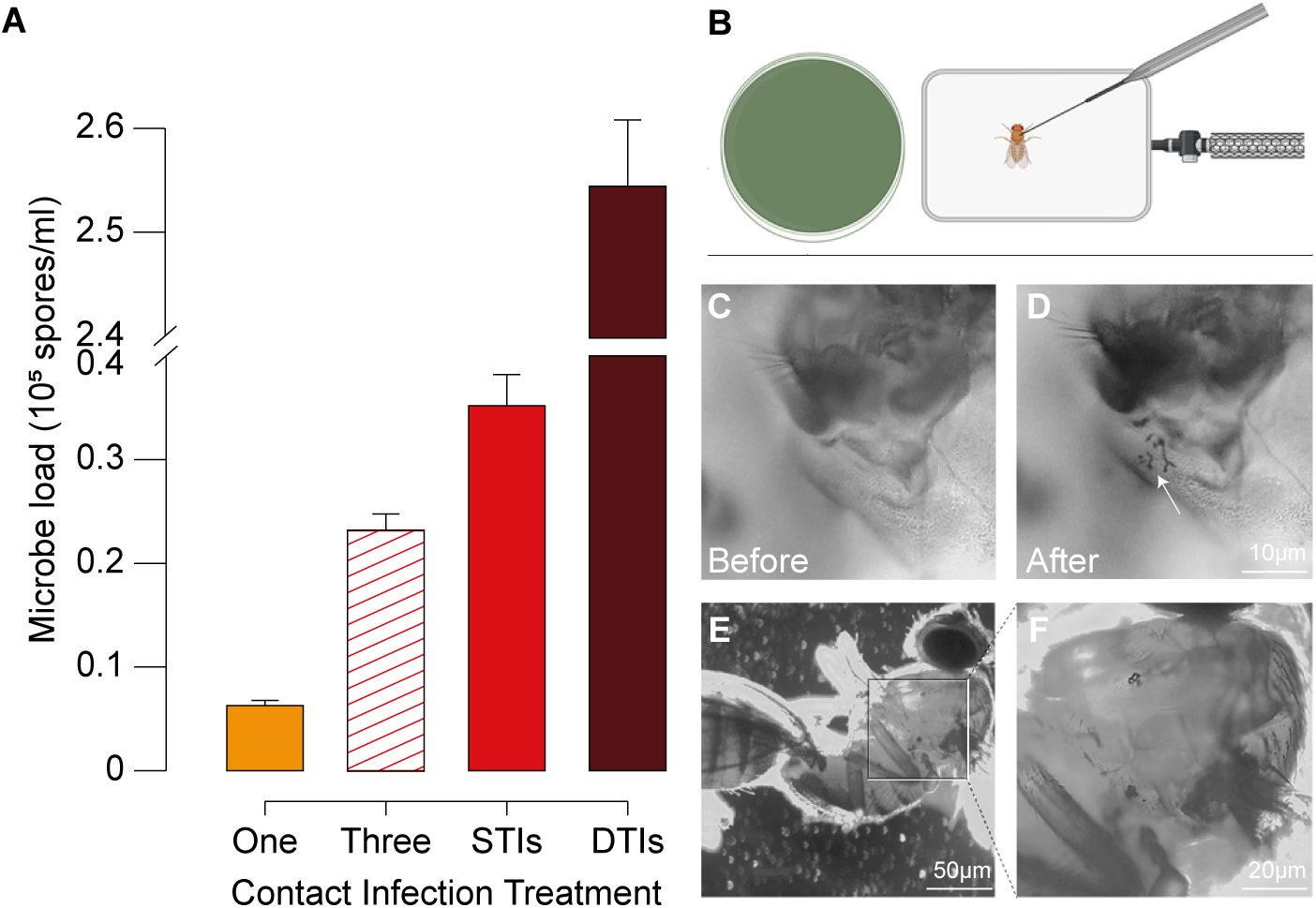
Modes of contact transmission. A, Inoculation loads of *Aspergillus austwickii* are reported for females treated with extremely low-dose (One Dose, yellow), low-dose (Three Doses, red slash), sexually transmitted infections (STIs, red), and direct topical infections (DTIs, brown) treatments (means ± SE). B, An illustration of the needle-tap inoculation method, which involves lightly tapping the Drosophila mesonotum once (One Dose) or three separate times (Three Doses) with a spore-covered tungsten needle. The duration of CO_2_ anaesthesia, number of inoculation taps, and the mating interval is controlled. The level of inoculation is visually confirmed for each batch. Fungal spore adhesion is shown, C and D, before and after inoculation with One Dose (Scale bar: 10µm). E, A naïve female (left) flash frozen during sexual congress with a DTIs treatment male (right) (50 µm). F, Spore adhesion evident on a DTIs male (20 µm).

We find that our modes of topical infection alter reproductive output in a manner consistent with the dynamic threshold model of terminal investment (Duffield et al., 2017). One Dose treatment demonstrates lower lifetime reproductive success (LRS) compared to Zero Doses and Three Doses treatments (*F*_(1,159)_ = 6.30, *p* = 0.013; F_(1,160)_ = 6.36, *p* = 0.013, respectively; Figure 2A). A single tap of live conidia reduces LRS by 17.0%, while increasing inoculation with two additional taps reverses this trend, leading to a 22.2% increase in LRS, comparable to the sham infection control, Zero Doses A (*F*_(1,153)_ = 0.02, *p* = 0.88). Similarly, STIs, with an inoculation dose akin to the three-tap treatment, yield consistent results, with increased inoculation resulting in LRS equivalent to that of the sham infection control (*F*_(1,168)_ = 0.19, *p* = 0.67). In contrast, the high-dose DTIs treatment significantly reduces reproductive output (*F*_(1,166)_ = 20.27, *p* < 0.001; Figure 2A).

**Figure 2.**
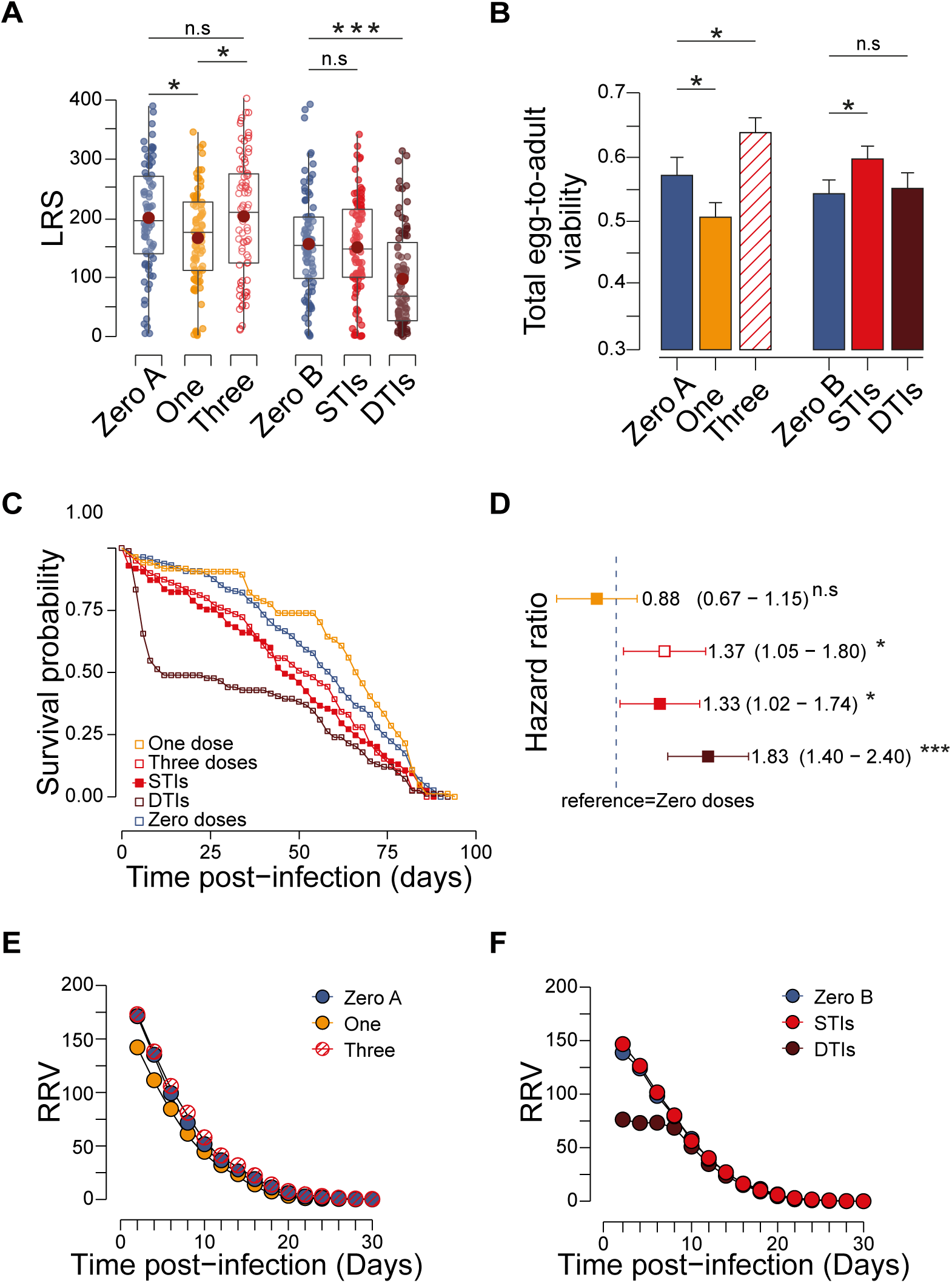
Terminal investment in Three Doses and STI treatments. **A**, Lifetime reproductive success (LRS, offspring numbers) is reported for Zero Doses A & Zero Doses B, blue), One Dose (yellow), Three Doses (red circles), STIs (red dots), and DTIs (brown) treatments (means ± SE; dots represent individual animals). **B**, Total egg-to-adult viability for each treatment (as before). **C**, Survival probability, and **D** hazard ratios (HR) is reported for each treatment (naïve treatments are pooled; HR is reported with 95% CIs). Statistical significance is reported as follows: * *p* < 0.05, *** *p* < 0.001, n.s not significant). **E** and **F**, Residual Reproductive Value (RRV) estimated for naïve (Zero Doses A & B, blue), One Dose (yellow), Three Doses (red slash), STIs (red solid) and DTIs (brown). See also Figure S1.

To determine whether the Three Doses and STI treatments induce terminal investment, we examined evidence of increased egg-to-adult viability alongside decreased post-infection survival. Although there is no significant variation in total egg number (*F*_(1,155)_ = 1.80, *p* = 0.18; *F*_(1,166)_ = 0.09, *p* = 0.76, respectively; Figure S1A), both the Three Doses and STI treatments lead to increased total egg-to-adult viability in female flies (*F*_(1,153)_ = 4.03, *p* = 0.047; *F*_(1,167)_ = 4.21, *p* = 0.042, respectively; Figure 2B). Notably, total egg-to-adult viability is calculated based on fertilised eggs capable of hatching. Despite ceasing to produce hatchable offspring within a month of starting reproduction, this viability is unrelated to the number of eggs laid during that period (Figure S1B). Concurrently, substantial reductions in survival rates are observed in the Three Doses and STI treatments (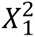 = 5.6, *p* = 0.018; 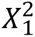 = 5.5, *p* = 0.019, respectively; Figure 2C), with significantly higher hazard ratios (HR = 1.37, *p* = 0.021; HR = 1.33, *p* = 0.032, respectively; Figure 2D). In contrast, the One Dose treatment reduces total egg-to-adult viability (*F*_(1,159)_ = 4.00, *p* = 0.047), yet shows the highest survival and the lowest hazard ratio among all treatments, although these differences are not statistically significant (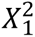 = 1.30, *p* = 0.30 and HR = 0.88, *p* = 0.34, respectively; Figure 2C and 2D). The DTIs treatment shows no effect on total egg-to-adult viability (*F*_(1,166)_ = 0.08, *p* = 0.78; Figure 2B), but exhibits substantially lower survival and a higher hazard ratio (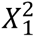 = 18.6, *p* < 0.001; HR = 1.83, *p* < 0.001).

### Terminal investment is triggered by instantaneous trade-offs rather than RRV

We find no support for the evolutionary explanation of terminal investment, which predicts that increased current reproductive success occurs when there are declines in future reproductive potential, specifically reduced RRV. Our findings indicate that for both the Three Doses and STI treatments, the RRV is higher, not lower, compared to females receiving the Zero Doses treatment (Figure 2E, 2F). Additionally, while there is evidence of reduced RRVs at early ages in the One Dose and DTIs treatments, both treatments also result in diminished reproductive output.

To test the alternative hypothesis that terminal investment results from synchronous trade-offs between reproduction and survival, we first determine the timing of peak egg-to-adult viability and then evaluate the subsequent changes in reproduction and survival over time. Overall, egg-to-adult viability varies with age across all infection treatments (*F*_(5,5701)_ = 28.59, *p* < 0.001, Figure 3A to 3D). As expected for an entomopathogenic fungus that takes several days to penetrate the insect cuticle and proliferate to cause infection (Hong et al., 2023), the first evidence of increased egg-to-adult viability is observed on days 6 and 8 post-infection for the Three Doses and STI treatments (*F*_(1,170)_ = 1.25, *p* = 0.007; *F*_(1,170)_ = 1.25, *p* = 0.011, respectively). Further analysis using age-specific models on both reproduction and mortality demonstrates that the benefits of infection to egg-to-adult viability decline concurrently with reductions in the cost of infection on mortality (repeated measures LME on viability: *t*_(1486)_ = −3.30, *p* = 0.001; *t*_(1375)_ = −2.36, *p* = 0.018, respectively; piecewise linear regression on mortality from day 8 to day 80: *t*_(34)_ = −2.25, *p* = 0.031; *t*_(34)_ = −2.28, *p* = 0.029, respectively, Figure 3B, 3D, 3E, Table S1, S2). The parameters estimated from these models indicate that the effects of infection on egg-to-adult viability and the natural logarithm of mortality rate diminish over time by 3.79% and 1.38% per day in the Three Doses treatment, and by 4.21% and 1.48% per day, respectively, in the STI treatment, thus supporting the synchronous trade-offs hypothesis for terminal investment. The age-dependent pattern observed at the lowest dose indicates that the impact of the infection on reproduction is immediate, with the decline in egg production due to fungal invasion diminishing over time (*t*_(1919)_ = 2.32, *p* = 0.021, Figure 3A, Table S1). In contrast, the effects of high-dose infections on the host remain consistent over time, showing no significant variation (*t*_(1293)_ = 0.92, *p* = 0.36, Figure 3C, Table S1).

**Figure 3.**
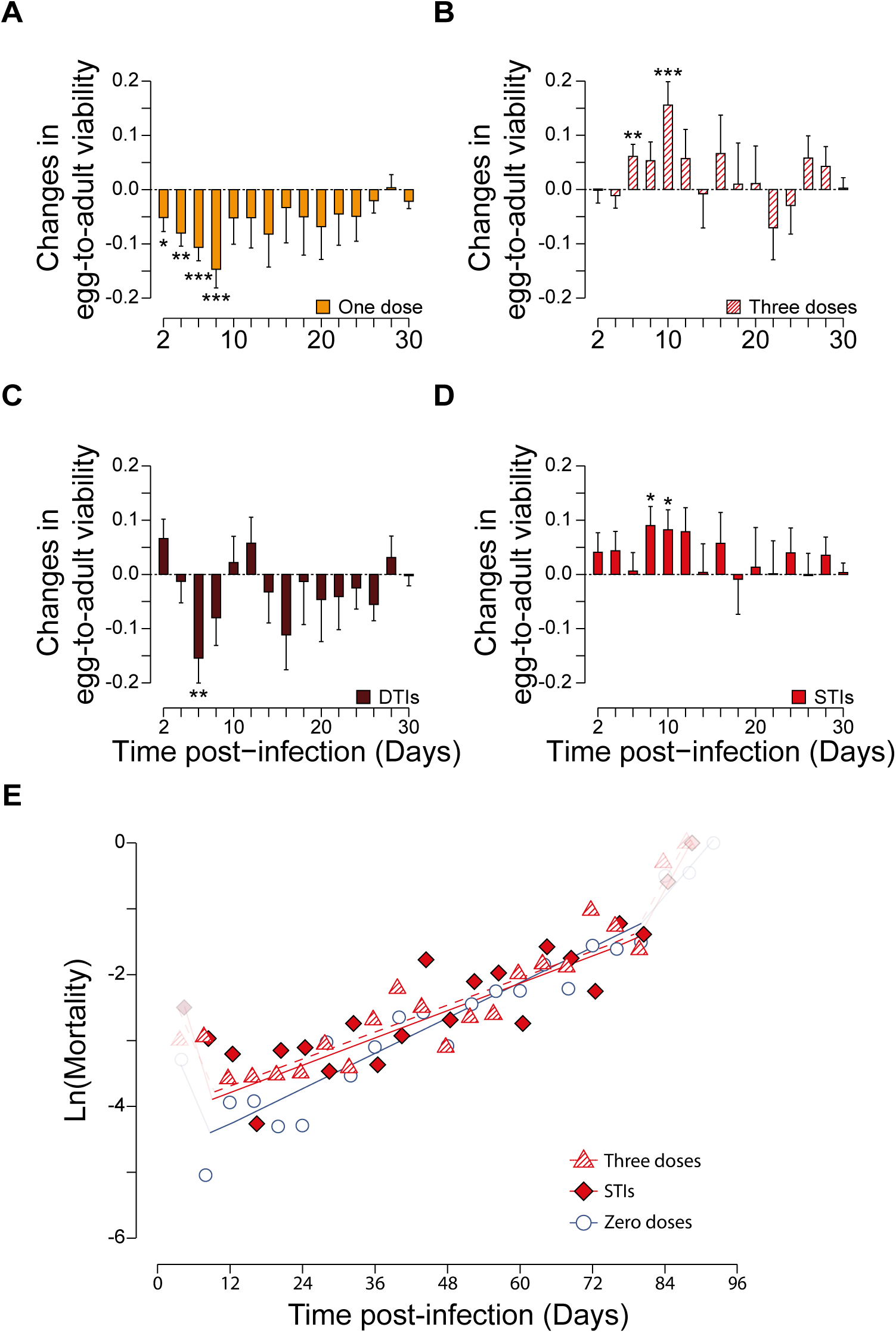
Synchronous trade-offs underlying terminal investment. **A** to **D,** Age-specific egg-to-adult viability of One Dose (yellow), Three Doses (red slash), DTIs (brown) and STIs (red) relative to the sham-infection naïve treatments (Changes in mean ± SE were estimated by subtracting values of the sham infection control from the infection treatment). Statistical significance is reported as follows: * *p* < 0.05, ** *p* < 0.01, *** *p* < 0.001. **E**, A piecewise linear regression of natural logarithm of mortality rate with age is reported for the critical interval from day 8 to day 80, comparing naïve (blue circle), Three doses (red slash triangle), and STIs (red rhombus) treatments. See also Table S1 & S2.

### Plastic life-history trade-offs depend on infection intensity

We find that the severity of infection alters how life-history traits trade-offs. Sham-infected organisms exhibit a strong negative correlation between fecundity and lifespan: higher total egg-to-adult viability leads to earlier death (r_(159)_ = −0.29, *p* < 0.001, Table S3). Low-dose infections maintain this negative correlation between traits, while increased inoculation doses intensify the strength of trade-offs (One Dose vs. Three Doses: Pillai = 0.12, *F*_(1,160)_ = 10.72, *p* < 0.001; STIs vs. DTIs: Pillai = 0.10, *F*_(1,165)_ = 9.51, *p* = 0.001; Zero Doses: r_(159)_ = −0.29, *p* < 0.001; One Dose: r_(82)_ = −0.37, *p* = 0.006; Three Doses: r_(76)_ = −0.47, *p* < 0.001; STIs: r_(82)_ = −0.41, *p* = 0.001, Figure 4, Table S3). In contrast, high-dose infections alter the relationship between the two traits (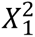 = 6.89, *p* = 0.032), resulting in no clear trade-off between total egg-to-adult viability and longevity (r_(81)_ = −0.17, *p* = 0.12, Figure 4, Table S3). Comparison with the Zero Doses control indicates that DTIs enhance the impact of longevity on fecundity. Females surviving DTIs exhibit significantly higher than expected total egg-to-adult viability (Generalised Linear Regression: Longevity × Infection: *t*_(160)_ = 2.16, *p* = 0.033, Table S3). However, low-dose infections do not alter the effect of individual longevity on fecundity (*t*_(156)_ = 1.25, *p* = 0.22; *t*_(150)_ = 1.42, *p* = 0.16; *t*_(161)_ = 1.08, *p* = 0.94; in the comparisons of One Dose, Three Doses, and STIs with their respective controls, Table S3).

**Figure 4.**
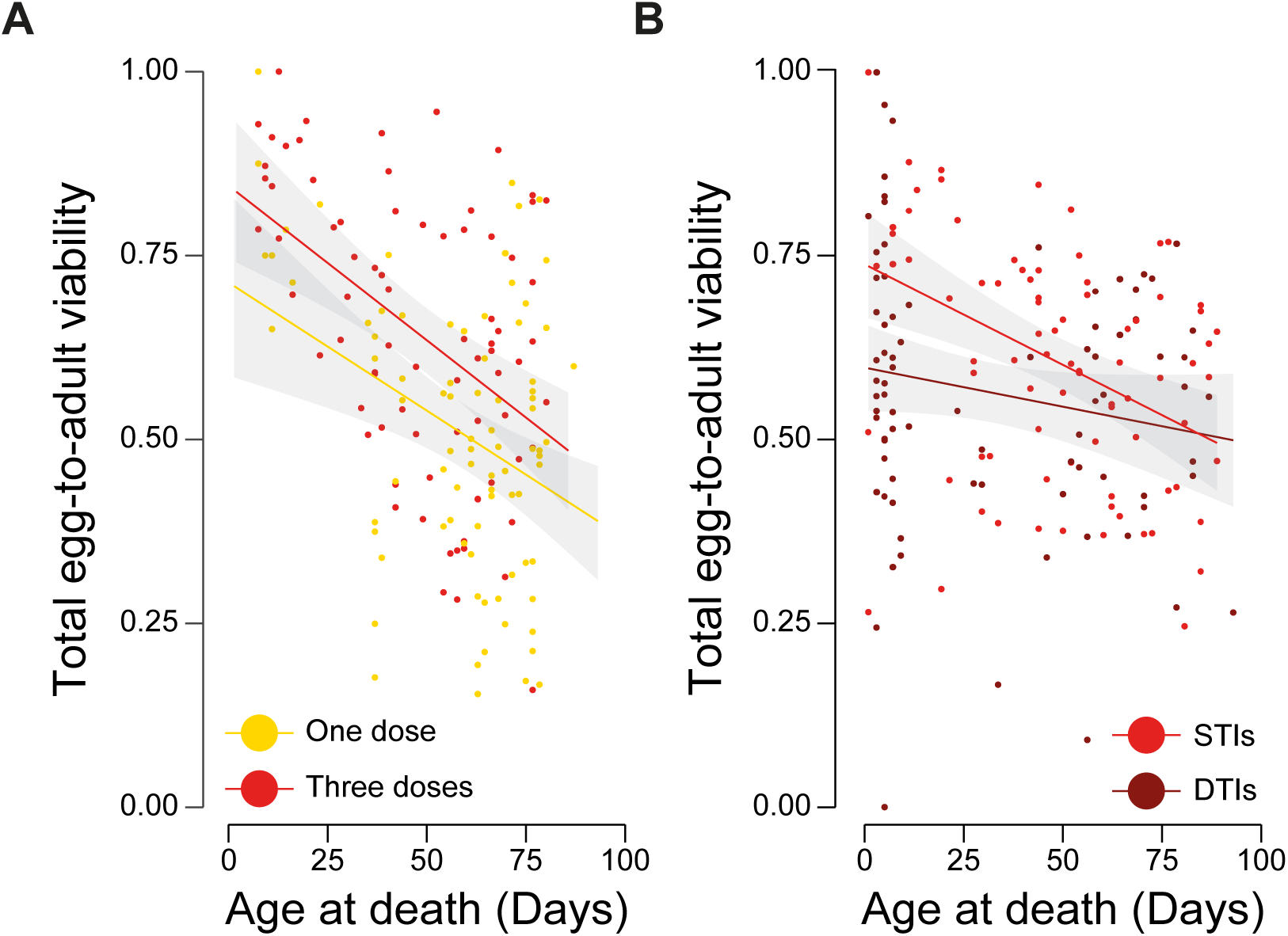
Infection severity as a determinant of life-history plasticity. **A**, Relationship between total egg-to-adult viability and age at death for individuals in One Dose (yellow) and Three Doses (red) treatments (95% CI represents fitting curves of each treatment, with dots indicating individuals). **B**, As before for STIs (red) and DTIs (brown) treatments. See also Table S3.

### The expression of *Turandot C* mediates terminal investment

We employed a candidate immune gene approach alongside large-scale biodemographic analysis to investigate the genetic basis of terminal investment. The *Turandot* genes in *D. melanogaster* encode a family of small peptides that play a role in the immune response, particularly when activated under stress and pathogenic challenge (An et al., 2012). Their notable non-immunological defence associated with mating behaviour suggests they may modulate reproductive responses to infection (see Immonen and Ritchie, 2012; Zhong et al., 2013). Our findings indicate that *TotC* expression provides early-age reproductive benefits following exposure to a model entomopathogenic fungus, *M. robertsii*, via STIs without incurring significant fitness costs. Age-specific analysis reveals that the impact of *TotC* expression on reproduction following STIs diminishes over time (Genotype × Time × Treatment: *t*_(932)_ = −1.97, *p* = 0.049, Figure 5B, Table S4). Specifically, wild-type expression of *TotC* (genotypes: +/UAS-*TotC*-IR & Act5C-Gal4/+) enhances offspring production on day one post-infection, while the line lacking *TotC* (Act5C-Gal4/UAS-*TotC*-IR) shows reduced fecundity (Genotype × Treatment: *F*_(1,154)_ = 9.11, *p* = 0.003; *F*_(1,151)_ = 10.53, *p* = 0.001, relative to each control line respectively, Figure 5A). In contrast, we examined the influence of *Dif*, the principal regulator of the Toll signalling pathway known for conferring immunity against fungal infections (Lemaitre et al., 1996), on reproductive outcomes. Our results demonstrate that *Dif* expression does not result in a fecundity boost on day one post-infection (Genotype × Treatment: *F*_(1,159)_ = 0.90, *p* = 0.35; *F*_(1,156)_ = 0.34, *p* = 0.56, comparing the knockdown line with each corresponding wild-type line, Figure 5C). Consistently, linear contrasts reveal that *TotC* expression has a significantly greater impact on terminal investment than *Dif* (*t*_(465)_ = −2.02, *p* = 0.044, Table S5).

**Figure 5.**
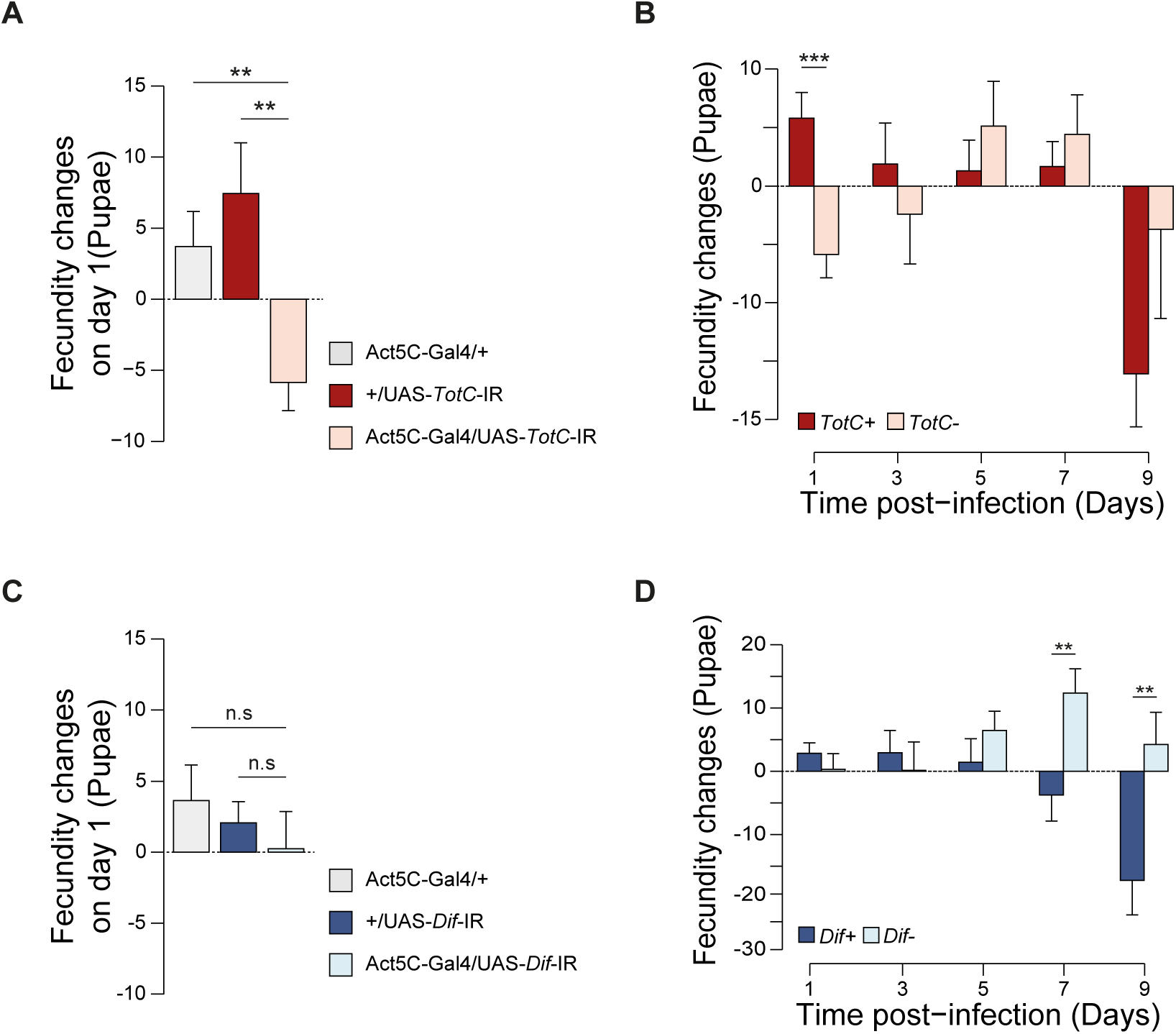
*TotC* rather than *Dif* expression promotes terminal investment. **A** and **C**, Changes in pupae numbers on day one following STIs of the two control genotypes (Act5C-Gal4/+, light grey; +/UAS-*TotC*-IR, rose red; and +/UAS-*Dif*-IR, blue), and the knockdown lines (Act5C-Gal4/UAS-*TotC*-IR, light pink; and Act5C-Gal4/UAS-*Dif*-IR, light blue) respectively. **B** and **D**, Changes in offspring numbers of the combined control genotype (Act5C-Gal4/+ & +/UAS-*TotC* -IR, rose red; Act5C-Gal4/+ & +/UAS-*Dif* -IR, blue, shown as *TotC+* and *Dif+*, respectively) and the knockdown line (Act5c-Gal4/UAS-*TotC*-IR and Act5c-Gal4/UAS-*Dif*-IR; shown as *TotC-*, light pink, and *Dif-*, light blue, respectively) across time. Changes in means ± SE were obtained by subtracting the number of pupae in the infection group from that in the sham-infection control. Significance is reported as follows: ** *p* < 0.01, *** *p* < 0.001, n.s, not significant. See also Table S4.

The reproductive advantages conferred by *TotC* expression, however, do not appear to incur any discernible costs. Both *TotC* knockdown and control lines exhibit neutral effects on longevity and fecundity following STIs (Genotype × Treatment: *F*_(1,293)_ = 0.05, *p* = 0.82; *F*_(1,230)_ = 0.05, *p* = 0.93, respectively, Figure 6A, S2A, S2B). Furthermore, the initial boost in reproductive output during early post-infection stages does not result in a loss of fecundity in the later reproductive phases tested (offspring produced between days 7 and 9 post-infection: Genotype × Treatment: *F*_(1,230)_ = 2.17, *p* = 0.14, Figure 5B). Conversely, the expression of *TotC* significantly decreases survival without corresponding reductions in reproduction under high-dose DTIs (Genotype × Treatment: *F*_(1,2448)_ = 23.36, *p* < 0.001; *F*_(1,260)_ = 1.32, *p* = 0.25, respectively, Figure 6B, S2C, S2D). This suggests that *TotC* expression facilitates fecundity compensation under STIs, but reduces survival under DTIs.

**Figure 6.**
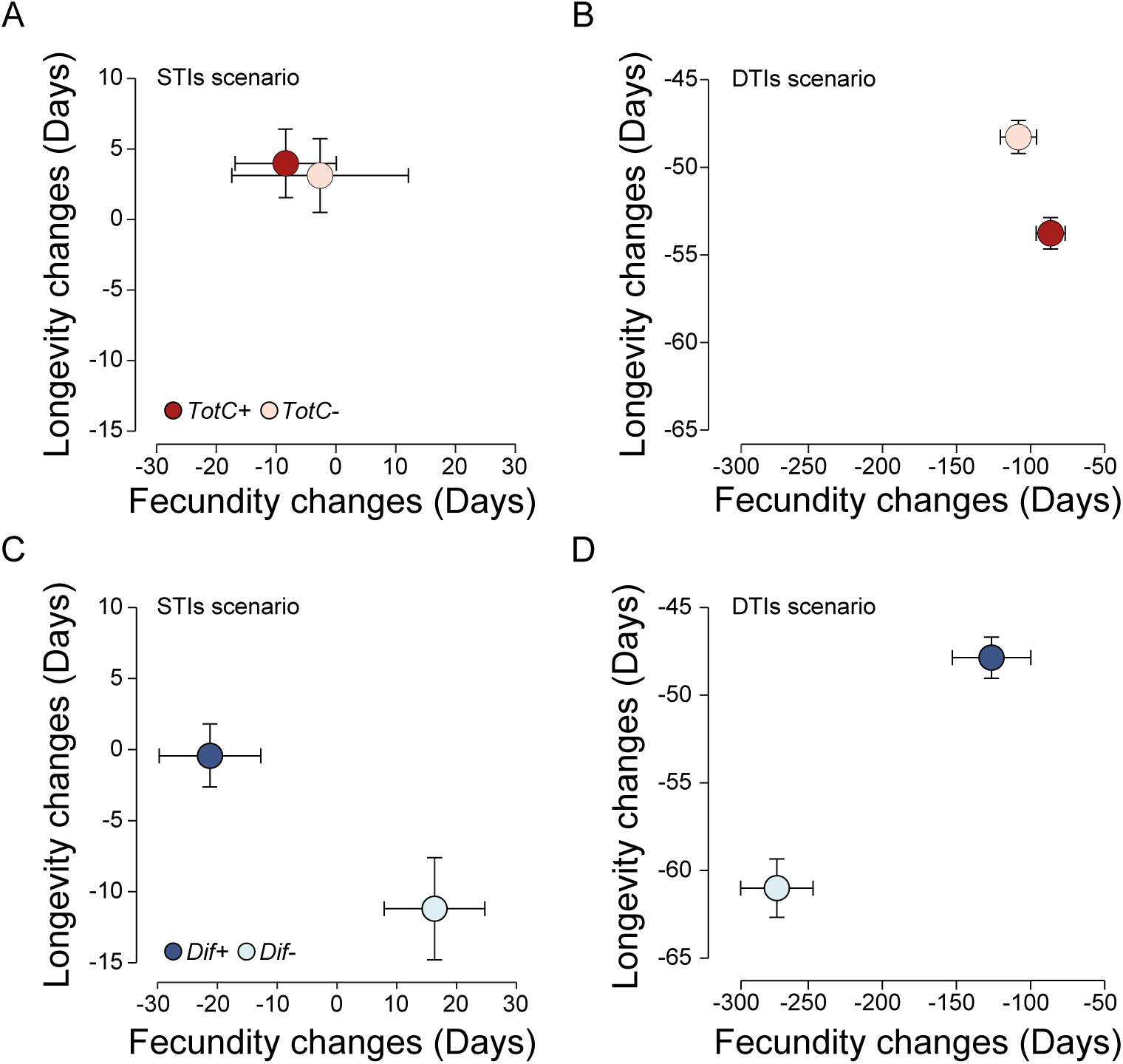
*Dif* expression, but not *TotC*, mediates antagonistic pleiotropy under STIs. Compared to sham-infected control, mean changes in longevity and fecundity due to *TotC* expression under **A**, STIs and **B**, DTIs. The combined control line (+/UAS-*TotC*-IR and Act5C-Gal4/+, shown as*TotC+*, rose red), the knockdown line (Act5C-Gal4/UAS-*TotC*-IR, shown as *TotC*-, light pink). **C** and **D**, as before for *Dif* expression under STIs and DTIs. Dots indicate the combined control line (+/UAS-*Dif*-IR and Act5C-Gal4/+, shown as *Dif+*, blue) and the knockdown line (Act5C-Gal4/UAS-*Dif*-IR, shown as *Dif*-, light blue). Changes in means ± SE were estimated by subtracting values of the sham-infection control from the infection treatment. See also Figure S2 & S3.

### *Dif-*mediated antagonistic pleiotropy under STIs

One of the crucial challenges to our discovery of low costs of TotC deployment to STIs is establishing how it differs from the effects of a canonical immune gene. We assessed the impacts of RNAi suppression of *Dif* on the reproduction and survival of female flies treated with *M. robertsii* following STIs and DTIs to determine whether the costs of *Dif* expression are higher than for *TotC*. Under wild-type expression of *Dif* (genotypes: +/UAS-*Dif*-IR & Act5C-Gal4/+; referred to as *Dif*+), we observe no significant difference in longevity between the STIs and naïve treatment groups (*F*_(1,215)_ = 0.04, *p* = 0.85). In contrast, *Dif*-females (knockdown genotype: Act5C-Gal4/UAS-*Dif*-IR) exhibits an average lifespan reduction of 11.9 ± 2.9 days (*F*_(1,102)_ = 17.58, *p* < 0.001, Figure S3A). However, a different pattern emerges concerning fecundity. We find genotype-specific differences in the effects of STIs on age-specific fecundity, with greater reductions observed in *Dif*+ females compared to *Dif*-females five days post-infection (LME, Genotype × Time × Treatment: *t*_(952)_ = −2.52, *p* = 0.012, Figure 5D, Table S4). Following infection, *Dif* expression decreases late-age (days 6 to 9 post-infection) reproduction by 21.30 ± 8.36 offspring (*t*_(144)_ = 2.55, *p* = 0.012), while the knockdown genotype results in a slight fecundity enhancement of 16.41 ± 8.17 compared to the sham infection control (*t*_(75)_ = −2.01, *p* = 0.048, Figure 5D, S3B). This indicates a pattern of terminal investment mediated by immune deficiency. The significant interaction between genotype and treatment affecting both longevity and late-age fecundity during STIs (*F*_(1,317)_ = 9.27, *p* = 0.003; *F*_(1,235)_ = 8.63, *p* = 0.004, respectively) suggests that *Dif* expression mediates antagonistic pleiotropy between reproduction and survival (Figure 6C). In contrast, we find that *Dif* expression enhances both traits under large-dose DTIs (Genotype × Treatment: *F*_(1,2695)_ = 138.45, *p* < 0.001; *F*_(1,251)_ = 46.29, *p* < 0.001, respectively, Figure 6D, S3C, S3D), showing no evidence of facilitating fitness trade-offs.

We conducted linear contrasts between *Dif* knockdown and control animals, comparing relative improvements in reproductive output for both infected and uninfected individuals across the STIs and DTIs treatments, to address the key question of whether *Dif* expression is more effective for reproduction under high-dose infections than low-dose infections. Our results show that *Dif* expression significantly increases the reproductive output of DTI-treated females compared to STI-treated animals (*t*_(486)_ = 5.97, *p* < 0.001, Figure S3E, Table S6).

## Discussion

The fact that we become ill suggests that evolving effective immune defences is inherently challenging. Studies of life history variation in infected insects are relevant to this issue, as they can reveal fundamental rules governing how animals cope under specific conditions and identify fitness trade-offs that impede the adaptive evolution of defence against infection. (McKean et al. 2008, McKean and Lazzaro 2011, Parker et al., 2011). Our investigation into life history trade-offs across a range of doses of topical fungal infections and host insect lines provides a surprising explanation for why immunity is often imperfect: insects have evolved to capitulate when faced with low-dose infections. Our findings indicate that terminal investment and innate immune responses are advantageous under low- and high-dose exposures, respectively, to the same entomopathogenic fungi, thereby reconciling the ecological contexts that drive the adaptive evolution of both non-immunological and immunological defence.

Life history theory posits that the evolution of immunity is accelerated by positive genetic correlations and constrained by negative genetic correlations between survival and fecundity (McKean and Lazzaro 2011; Rauw 2012; Schwenke et al. 2016). Increasingly, studies emphasise “modular pleiotropy,” wherein genes influence specific clusters of traits rather than broadly affecting the entire phenotype (Hansen 2003; Stearns 2010; Houle and Rossoni 2022). In our Gal4 lines, *Dif*-mediated phenotypic plasticity affects only offspring number, not multiple reproductive subtraits (e.g., egg-to-adult viability, as observed in our Dahomey lines). We find no evidence of fitness or immunological trade-offs in the high-dose treatments, suggesting that mechanisms improving survival can evolve freely when the infectious dose is sufficiently high. However, under STIs, the expression of *Dif* mediates strong antagonistic pleiotropy and generates negative correlations between survival and reproduction, indicating that under common infections, the adaptive evolution of immunity is constrained by the coupling of immune function and egg production (Zerofsky et al. 2005; He and Zhang 2006; Van Rheenen et al. 2019).

Although terminal investment induced by parasitic challenges is regarded as an adaptive non-immunological mechanism, it remains unclear how succumbing to infection mitigates fitness costs associated with immunological responses (Minchella and Loverde 1981; Polak and Starmer 1998; Bonneaud et al. 2004; Weil et al. 2006; Parker 2011; Brannelly et al. 2016; Hudson et al. 2020; Schulz et al. 2023; Liao et al. 2024). In our wild-type lines, the net effect of terminal investment is neutral; hosts inoculated with low doses of conidia (Three Doses and STI treatments) do exhibit reduced survival and lower net egg production; however, they achieve equivalent lifetime reproductive success (LRS) compared to sham-treated controls due to increases in egg-to-adult viability occurring during periods of heightened mortality. In the test of *Dif* expression’s consequences, evidence indicates that it inhibits terminal investment under low-dose infection, as STI-treated females show higher LRS but reduced longevity compared to controls in the absence of *Dif* expression. While, in *TotC* lines, *TotC* expression promotes fecundity compensation in STI-treated females by enhancing early-age reproduction, without incurring costs to LRS or longevity. These findings suggest that *TotC* facilitates fecundity compensation without evident fitness costs, while *Dif* enhances post-infection survival at substantial fitness costs.

The ability of *Turandot* family genes to mitigate cytotoxicity accounts for the low costs associated with *TotC* expression. As a central component of the Toll signalling pathway, *Dif* provides effective protection against pathogenic fungi and Gram-positive bacteria by regulating the expression of target genes, including antimicrobial peptides (AMPs), which can increase cytotoxicity to host cells (Ip et al., 1993; Lemaitre et al., 1996; Bacalum and Radu, 2015). In contrast, recent studies have demonstrated that *Turandot* family proteins protect host tissues from damage inflicted by cationic pore-forming AMPs (Rommelaere et al., 2024). The *Turandot* gene family also plays broader roles in stress responses to infection, heat shock, and radiation (Ekengren and Hultmark, 2001; Agaisse et al., 2003; Amstrup et al., 2022). Additionally, *TotC* expression is upregulated in response to mating cues, suggesting its involvement in immune anticipation and other processes that may limit immunological costs (Immonen and Ritchie, 2012; Zhong et al., 2013). Combined with previous findings indicating that *TotC* does not influence microbe load (Zhong et al., 2013), our study illustrates that *TotC*-regulated terminal investment serves as a mechanism of non-immunological defence that promotes tolerance rather than resistance to infection (Schneider and Ayres, 2008; Rafaluk-Mohr et al., 2022). These findings are consistent with previous research demonstrating mechanisms of capitulation in response to low-dose STIs (Adamo, 2014).

The central assumption of the terminal investment hypothesis is that terminal occurs because current reproduction trades off against future reproductive potential, explicitly defined as Residual Reproductive Value (RRV) (Williams, 1966; Hirshfield and Tinkle, 1975; Pianka and Park, 1975; Clutton-Brock, 1984; Duffield et al., 2017; Stearns, 1992). We find no support for this assumption. Employing the method of Pianka and Park (1975), we discover that RRV is higher, not lower, in the Three Doses and STI treatments, which exhibit terminal investment. Conversely, RRV is substantially lower in the DTIs treatment, where, according to theory, it should have been higher. Thus, the terminal investment phenomenon we report here does not reflect a lifelong “terminal” shift in reproductive strategy.

Though asynchronous trade-offs between early reproduction and late-life survival drive are considered to be core drivers of life history evolution, there has been less attention on the importance of synchronous trade-offs (Williams, 1957; Kirkwood and Rose, 1991; Lemaître et al., 2015 & 2024). Our study shows that terminal investment is caused by synchronous trade-offs. Using piecewise regression, we find that increases in egg-to-adult viability occur concurrently with rises in mortality rate during the first 10 days post-exposure to low-dose infections. In the wild-type line, we observe strong negative correlations between egg-to-adult viability and longevity, suggesting that individuals with generally higher reproductive effort tend to have shorter lifespans, and vice versa. Notably, a similar lack of association between ageing and reproductive value has also been demonstrated in rockfish (Kolora et al., 2021).

Previous research indicates that both infectious dose and mode of infection influence the severity of infection and the dynamics of host-pathogen coevolution (Ebert and Mangin, 1997; Zhong et al., 2013; Martins et al., 2013). Our study contributes to these findings by demonstrating that the mode of topical infection determines whether hosts employ mechanisms to combat or capitulate to infection, potentially selecting for increased microbial virulence or “mafia-like” parasite manipulations (Thomas et al., 2005; Parker et al., 2011). The majority of infection research relies on septic infections induced via artificial injection or body puncture, which can obscure natural processes and often result in rapid mortality, preventing any reproductive responses to infection (Wigby et al., 2008; Lu and St. Leger, 2016; Dong and Dimopoulos, 2023). Current research methodologies frequently overlook the capacity of organisms to anticipate, detect, and avoid infections, which are likely critical non-immunological defence mechanisms (Park et al., 2011; Ortiz-Urquiza and Keyhani, 2013).

## Materials and methods

### Fly strains and fungal culture maintenance

We paired different strains of flies and fungi for various aspects of our study. To maximise the ecological relevance of our research on life history trade-offs associated with terminal investment, we aimed to simulate the pathogens and common infection patterns frequently encountered by *Drosophila* populations in nature. For this purpose, we utilised the Dahomey stock as a host for *Aspergillus austwickii*, an entomopathogenic fungus isolated from and endemic to the Dahomey population. Additionally, to investigate the genetic mechanisms underlying post-infection fitness trade-offs, we employed recently isogenized lines from the UAS/Gal4 gene expression system as hosts for the model entomopathogenic fungus *Metarhizium robertsii*.

*A. austwickii* was cultured from a single colony obtained 12 months prior to the start of the experiments from the Dahomey population of D. melanogaster (gratefully provided by Prof. Tracey Chapman, University of East Anglia). The fungal colony was derived from a 4-day-old mated female fruit fly that had been anaesthetised, surface-sterilised, homogenised, and plated at a 1:100 dilution on standard malt extract agar (MEA). Notably, this fungus was recoverable from all tested females (20/20) from the Dahomey stock. *M. robertsii* was cultured from spores obtained from the Agricultural Research Service Collection of Entomopathogenic Fungal Cultures (isolate 2575, ARSEF, United States Department of Agriculture). This strain has been extensively used in studies focusing on innate immunity in insects. This fungus is widespread in soil and exhibits high pathogenicity towards insects (Roberts and St Leger, 2004). Unlike experiments with *A. austwickii*, inoculating *Drosophila* with *M. robertsii* allows us to verify the cause of death, as it is not naturally present in the Dahomey stock. For spore collection, we established a standard 3-month fungal culture procedure by conducting trials at 25°C and 50% humidity on MEA plates, where spores were isolated from culture every month. The 3-month duration consistently yielded minimal evidence of spore-to-spore adhesion, as confirmed under light microscopy.

The fly stocks were maintained at 25°C and 50% humidity under a 12:12 light-dark photoperiod, using standard oatmeal–molasses–agar media fortified with an antifungal agent (Nipagin) and an acid mix containing propionic and phosphoric acid to inhibit bacterial growth, with a single grain of baker’s yeast supplemented on the surface. To control for ecological factors, the fly cultures were maintained for three generations in standard *Drosophila* vials at medium density (approximately 40-60 flies per vial) prior to the start of the experiments, as population density can affect fitness traits (Than et al., 2020).

The UAS-*Dif*-IR and UAS-*TotC*-IR strains were obtained from the Vienna *Drosophila* RNAi Centre. We chose the Act5C-Gal4 constitutive promoter (Act5C-Gal4/CyO, Bloomington Stock Centre, stock number 4414) to induce constitutive expression of Gal4, which we confirmed, through semi-quantitative PCR, reduces expression of both *Dif* and *TotC* when co-inherited with the UAS-*Dif*-IR and UAS-*TotC*-IR vectors, respectively. To minimise potential bias from mutation accumulation in the genetic background of the lines, we followed four procedures. First, we crossed each of the lines to a w1118 line (the genetic background for the RNAi strains) that had been recently isogenized for one generation prior to the start of the experiment. Second, we tested multiple control lines. To generate the two lines with reduced target gene expression (Act5C-Gal4/UAS-*Dif*-IR and Act5C-Gal4/UAS-*TotC*-IR), we crossed Act5C-Gal4/CyO females with males carrying the UAS construct. Control lines were established to account for the potential effects of Gal4 transcriptional activator expression and the presence of the UAS constructs by crossing the RNAi line females (Act5C-Gal4/CyO females) with w1118 males, resulting in three control lines: Act5C-Gal4/+, UAS-*Dif*-IR/+, and UAS-*TotC*-IR/+. Third, we developed a statistical pipeline that focuses on Gene × Infection interactions, rather than genetic effects within a single infection treatment, to ensure that the consequences of gene expression are influenced by the infection context. Finally, we plotted changes in the fitness consequences of infection within each line in comparison to sham controls to eliminate potential biases from line-specific mutations unrelated to changes in target gene expression.

### Fly mating and infection

We varied the topical infection load of female fruit flies using a range of methods. To coat the insect cuticle with an abundance of live *A. austwickii* spores, we employed the Direct Topical Infections (DTIs) method, often referred to as the “natural infection” method. This involved sexing virgin flies within 4 hours of emergence under light anaesthesia, mating them on day 2 over a 24-hour period, and infecting them on day 3 by gently swirling cohorts of 120 flies in a 50 ml conical flask containing 6 mg of mature fungal spores for 20 seconds without anaesthesia. To inoculate females with low-dose Sexually Transmitted Infections (STIs), we collected virgin flies as before, treated cohorts of male flies on day 2 with the DTIs treatment, placed them in temporary cages with fresh media (to facilitate self-grooming), and mated the virgin females on day 3 by transferring 30 DTI-treated males without anaesthesia to cages containing 30 virgin females for 12 hours.

Subsequently, individual females were transferred without anaesthesia to individual shell vials. The comparable Zero Dose control treatment involved the same procedure as the STI treatment, except that females were provided with males swirled in flasks for 20 seconds without mature fungal spores. We developed a needle tap method to inoculate females with low-dose topical infections, which involved mating females on day 2 over 24 hours and subsequently conducting three separate inoculations by running the tip of a tungsten needle through a lawn of mature fungal spores and lightly tapping the mesonotum of lightly anaesthetised flies (Three Doses Treatment). For administering an extremely low-dose topical infection, mated females were gently tapped once with live spores and twice with a sterile needle (One Dose Treatment). The comparable Zero Doses control treatment involved three gentle taps with a sterile needle (Zero Doses control). In each of the three needle tap treatments, the duration of inoculation (including sterile controls) was set at 100 seconds to minimise variation resulting from light anaesthesia.

### Microbe load measurement

We tested the consistency of the pathogen burden by counting fungal spores using a haemocytometer following the administration of Zero Doses, One Dose, Three Doses, STIs, and DTIs treatments. Cohorts of 10 treated females were placed into a 10 ml centrifuge tube containing 100 µl of 70% ethanol and mixed on a vortex mixer (1500 rpm for 10 seconds) to dislodge and suspend spores from the fly cuticle. A volume of 15 µl of spore suspension was then added to each chamber of the haemocytometer (10 chambers, 4×4 grid), applying dilutions for ease of counting: no dilution for the Zero Doses and One Dose treatments, 2-fold dilution for the Three Doses and STI treatments, and 10-fold dilution for the DTIs treatment. After allowing the suspension in each chamber to settle, spores were counted within five non-adjacent grids under a microscope. Three technical replicates were scored per sample, and this process was repeated three times. The concentration of spores per ml was calculated by multiplying the number counted per grid by the inverse of the sample dilution. Fungal loads across all treatment groups were normalised by subtracting the spore counts from the Zero Doses treatment group.

### Life history changes from topical inoculations

After the females were inoculated under the Zero Doses, One Dose, Three Doses, STIs, and DTIs treatments, dead flies were removed (∼1% of treated animals), and the surviving females were placed into assigned positions within randomised blocks (n = 90/treatment). To assess their reproductive success, the experimental subjects were transferred to fresh vials every 24 hours until day 8 post-infection, then every two days until day 30, and subsequently every four days until death. With each transfer to a fresh vial, the number of eggs was recorded. The number of pupae that hatched from those eggs was counted 17 days after the eggs were laid. Egg-to-adult viability was calculated as the ratio of pupae to eggs. Escaped or sterile individuals were excluded from data analysis, although pupa and egg-to-adult viability records were retained up to the day of escape. From these counts, we estimated total and age-specific measures of egg production, reproductive success, egg-to-adult viability, and longevity for females across each infection treatment.

### Residual Reproductive Value measurement

Our experiments were designed to test whether terminal investment arises from trade-offs between current and expected reproduction (supporting the widely held explanation that current reproductive effort increases when future reproductive output decreases) or from trade-offs between current reproduction and current hazard. We calculated Residual Reproductive Value (RRV) for each treatment using the method described by Pianka and Park (1975), where RRV is defined as an organism’s reproductive value in the next reproductive age multiplied by the ratio of next survivorship to current likelihood of survival, expressed as:

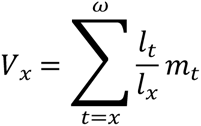

Here, *x* and *t* represent age, and *ω* indicates the age of last reproduction, while *m_t_* represents fecundity at age *t*. RRV can be delineated as follows:

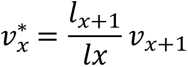

In this equation, *v_x_\** denotes the residual reproductive value, *l_x_* represents age-specific survivorship, and *v_x+1_* is the reproductive value in the next age interval. Based on the theoretical expectation that declines in RRV drive terminal investment, we would expect estimated RRV to decline relative to controls across each age group for treatment that generates terminal investment.

### Genetic trade-offs in RNAi lines

To evaluate the impact of *TotC* and *Dif* gene expression on life-history trade-offs in animals exposed to varying infection loads, we ran large-scale demographic studies to obtain estimates of lifetime reproductive success and adult longevity. Each of the transgenic lines was back-crossed to the control w1118 line to minimise potential effects from line-specific deleterious recessive mutations. After confirming that the knockdown effectively reduced gene expression relative to control lines, we measured the reproductive success and lifespan of the lines (Act5C-Gal4/UAS-transgene-IR) alongside their control lines (UAS-transgene-IR/+ and Act5C-Gal4/+), following treatment with STIs or DTIs (Zhong et al., 2013). The number of pupae was recorded daily until day 9 post-infection, and lifespan data were recorded every two days after day 9 until death.

### Statistics

Our statistical pipeline of the analysis of variation in reproduction was to conduct one-way ANOVA to confirm differences between treatments and lines, but rely on two-way ANOVA, to establish that changes in fitness traits depended on treatment contexts. Similarly, survival was assessed using the Kaplan-Meier model and Cox proportional hazards regression, with significant difference in main effects established first, followed by tests for context-specific change in fitness traits. Age-specific fecundity was evaluated using a linear mixed-effects model (LME, Pinheiro et al., 2012). Piecewise regression was conducted to analyse segmented age-dependent mortality rates (Table S2). Combined MANOVA (Pillai’s Trace test), Pearson correlation tests, and general linear regression were employed to evaluate the influence of infection dose on the correlation between lifespan and fecundity. We developed a statistical pipeline to establish evidence of genetic trade-offs in the RNAi lines. For each analysis of *TotC* and *Dif*, we first confirmed that gene expression in the knockdown lines generated significantly different fitness trait values compared to their associated transgene controls. To ensure that the observed genetic effects did not stem from line-specific deleterious recessive mutations, we focused on evidence of line × infection treatment statistical interactions and confirmed that the knockdown line was the source of the interaction by running analyses with and without the knockdown. Only after verifying that there were no significant differences between the two control genotypes (+/UAS-transgene-IR & Act5C-Gal4/+), did we proceed with analyses using the combined control genotype. A linear contrast model was utilised to assess the effectiveness variance between *TotC* and *Dif* genes on fecundity under STIs, along with the differential reproductive consequences mediated by *Dif* expression under varied infection modes (Table S5 & S6). A two-tailed Student’s t-test was conducted after performing an LME analysis to compare reproductive differences between genotypes. All statistical analyses were performed using R version 4.3.1, and p-values lower than 0.05 were deemed statistically significant.

## Author Contributions

JL and NKP conceived the research. JL and NKP conducted the experiments and data collection. JL analysed the data. JL wrote the manuscript. Both authors contributed critically to the drafts and gave final approval for publication.

## Acknowledgements

We thank Madeline Doyle and Jake Creasey for their assistance with data collection. This study was supported by a Royal Society Research Grant and a BBSRC, Defra, NERC, Scottish Government, and Wellcome Trust grant, BB/I000836/1, to NKP.

## Data availability statement

The data supporting the findings of this study can be found on Figshare via the identifier: https://figshare.com/s/f556eb28c0c1ecde5e8d

## Supplementary materials

**Figure S1.**
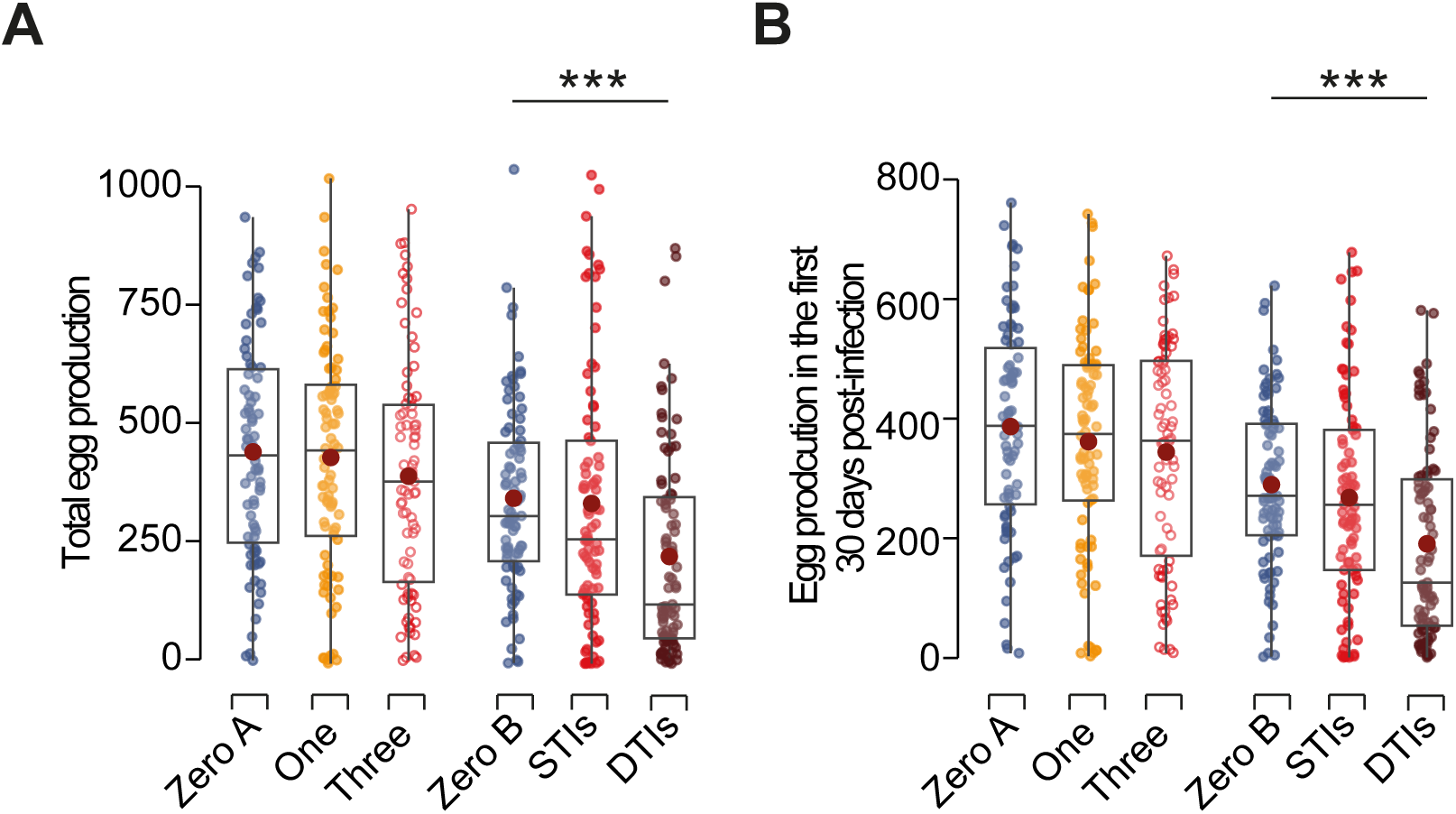
Terminal investment is not stimulated by an increase in egg production. Related to Figure 2. **A**, Total egg production, and **B,** Egg production within the 30 days post-infection, which is the period during which hatchable offspring are produced, is reported for the following treatments: Zero Doses A & B (blue), One Dose (yellow), Three Doses (red circles), STIs (red dots), and DTIs (brown). Means ± SE, and dots represent individual animals. Significance is reported as follows: *** *p* < 0.001. Note: nearly all zygotes are produced within 30 days following infection. Changes in egg-to-adult viability are not influenced by variations in egg production during this period, emphasising that terminal investment is driven by enhanced reproductive quality rather than quantity.

**Figure S2.**
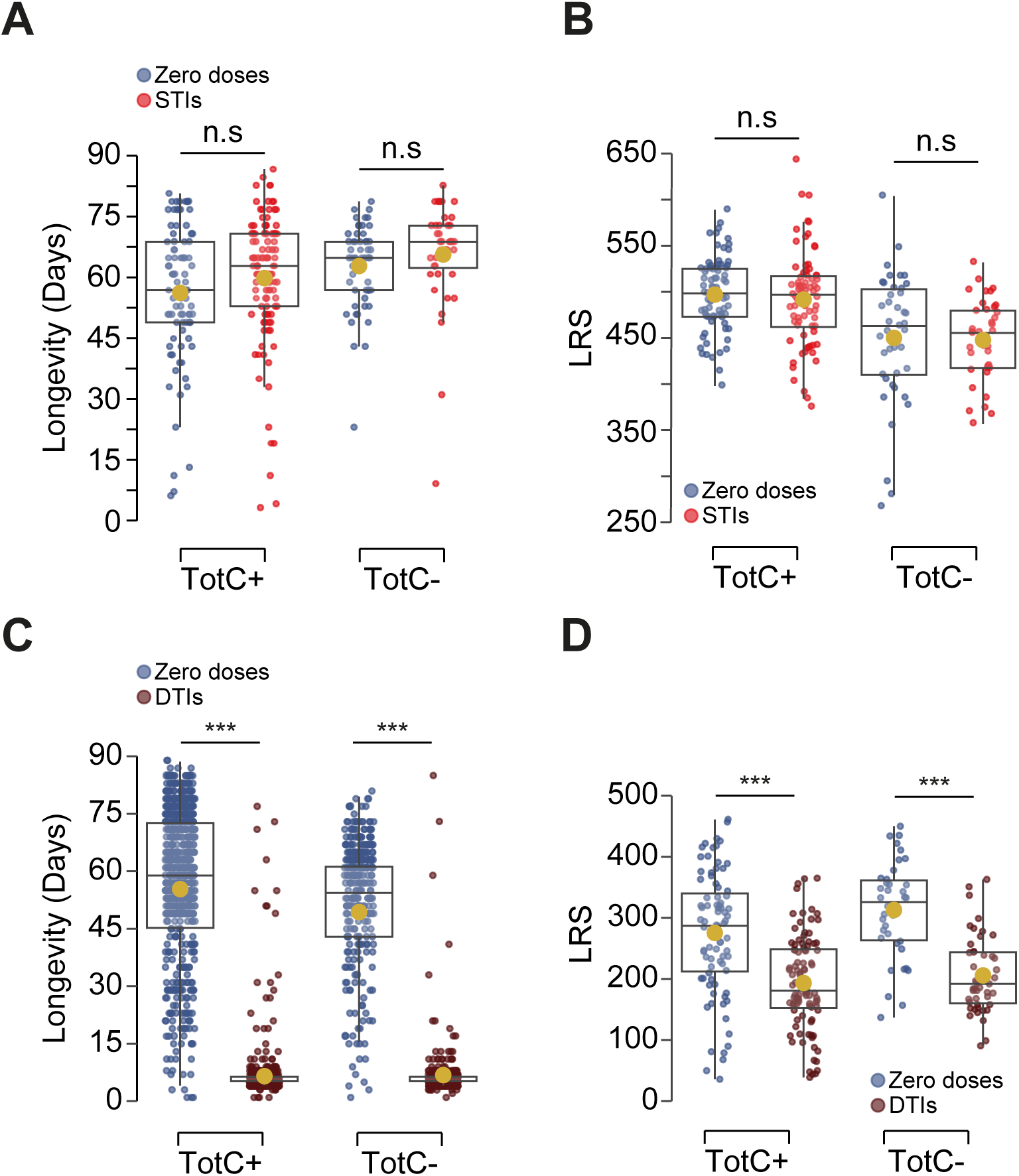
*TotC* expression triggered terminal investment shows inconspicuous fitness costs. Related to Figure 6. **A**, Longevity, and **B**, LRS of the combined control line (Act5C-Gal4/+ & +/UAS-*TotC*-IR, shown as *TotC+*) and the knockdown line (Act5c-Gal4/UAS-*TotC*-IR, shown as *TotC-*) are reported for the sham-infection naïve treatments (blue) and STIs (red). **C**, Longevity, and **D**, LRS of the combined control (*Dif*+) and the knockdown line (*Dif*-) under sham-infection treatment (blue) and DTIs (brown). Dots represent individual animals with a yellow dot indicating the mean. Significance is reported as follows: *** *p* < 0.001, n.s, not significant.

**Figure S3.**
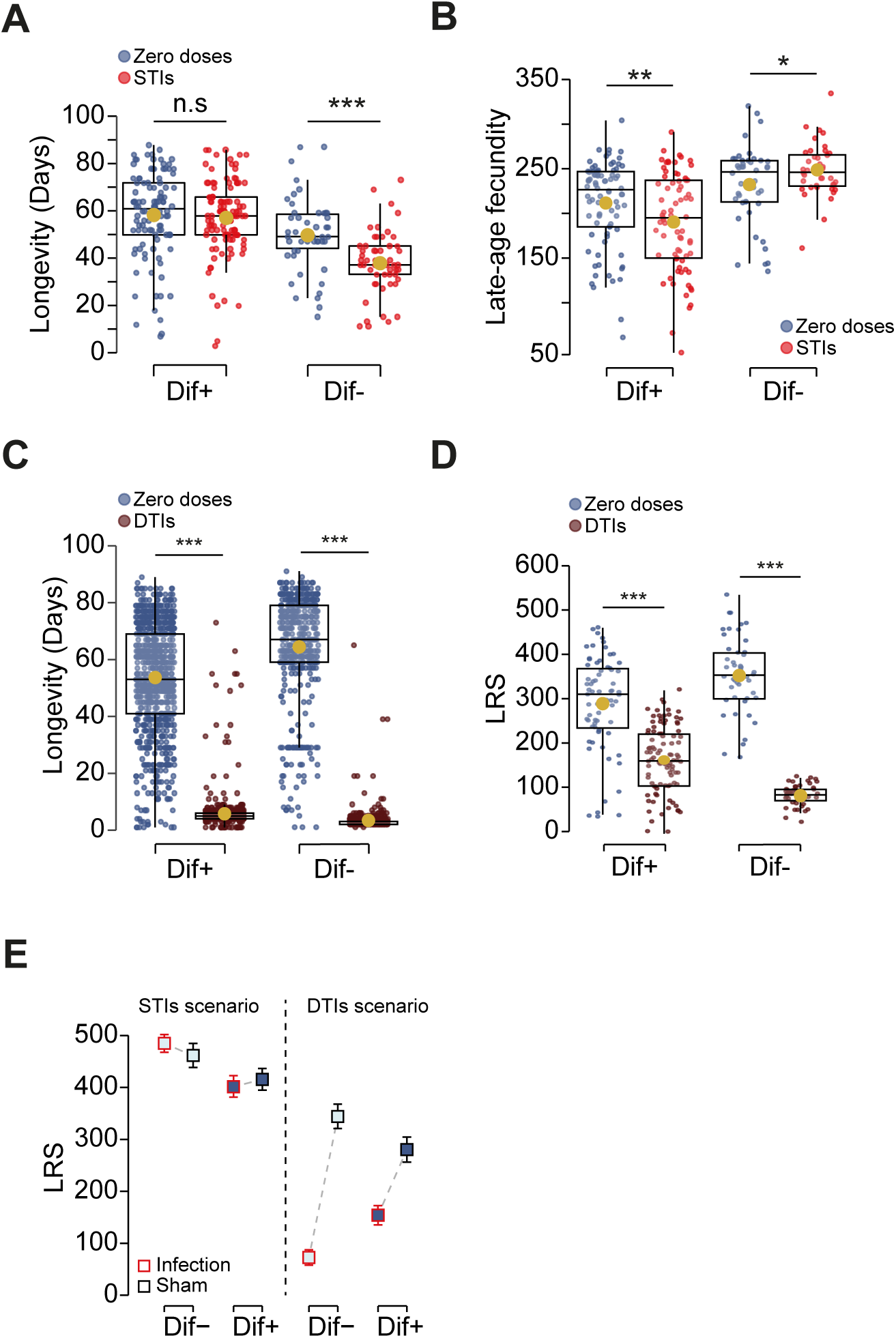
Trade-offs post-infection facilitated by *Dif* expression are dependent on mode of infection. Related to Figure 6. **A**, Longevity, and **B**, Late-age reproduction (day 6 to day 9) of the combined control line (Act5C-Gal4/+ & +/UAS-*Dif*-IR, shown as *Dif+*) and the knockdown line (Act5c-Gal4/UAS-*Dif*-IR, shown as *Dif-*) are reported for the sham-infection naïve treatments (blue) and STIs (red). **C**, Longevity, and **D**, LRS of the combined control (*Dif*+) and the knockdown line (*Dif*-) under sham-infection treatment (blue) and DTIs (brown). **E**, LRS of the combined control line (*Dif+*, blue) and the corresponding knockdown line (*Dif-*, light blue) following infection (red) and sham infection treatment (blue) under STIs and DTIs scenarios. The dashed lines indicate distance between each pair-comparison, representing a contrast of fitness changes under different modes of infection. Dots represent individual animals with a yellow dot indicating the mean. Significance is reported as follows: * *p* < 0.05, ** *p* < 0.01, *** *p* < 0.001, n.s, not significant.

**Table S1.**
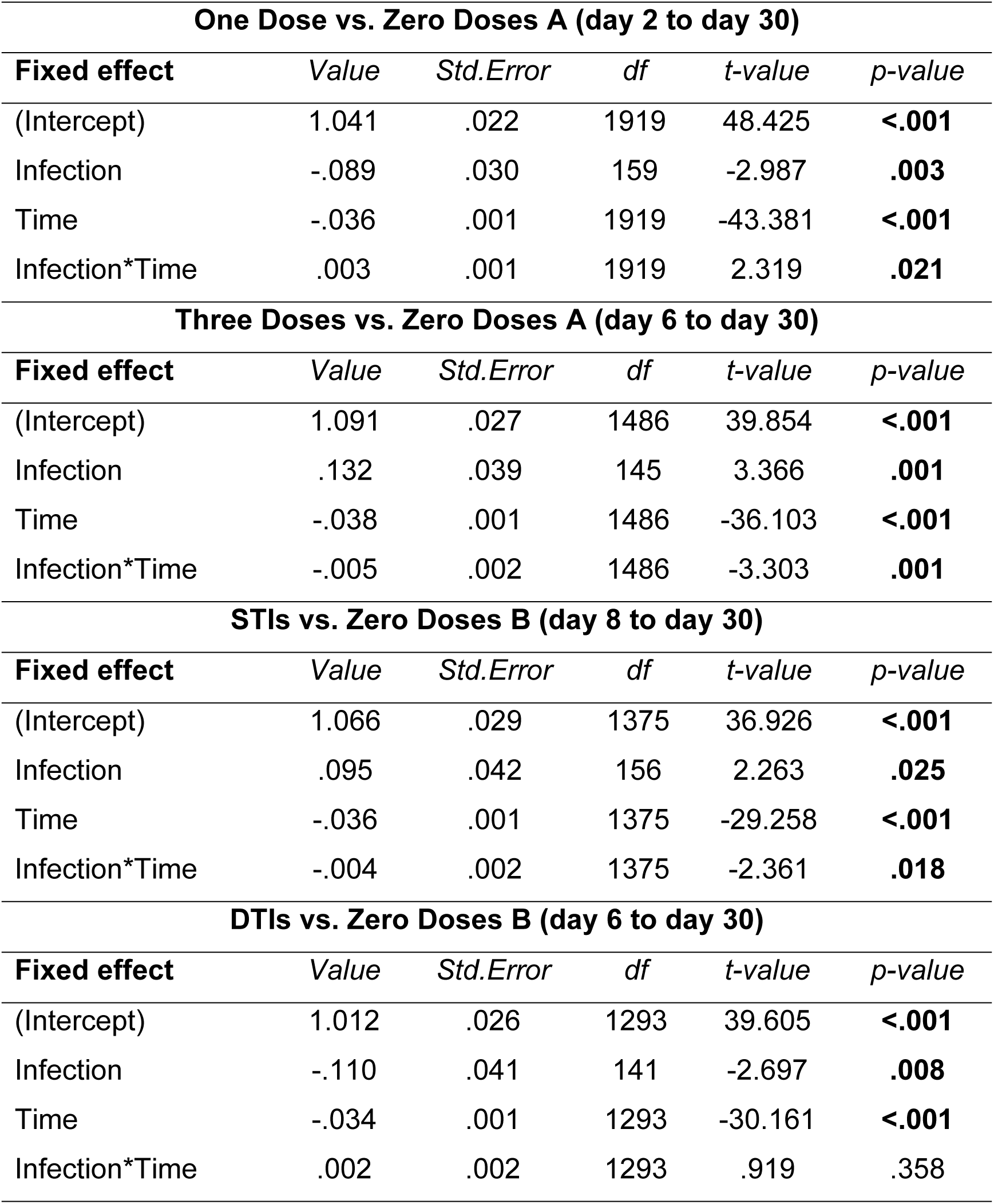
Results of LME on age-specific egg-to-adult viability. Related to Figure 3A to 3D.

| **One Dose vs. Zero Doses A (day 2 to day 30)** |  |  |  |  |  |
| --- | --- | --- | --- | --- | --- |
| **Fixed effect** | *Value* | *Std.Error* | *df* | *t-value* | *p-value* |
| (Intercept) | 1.041 | .022 | 1919 | 48.425 | **<.001** |
| Infection | -.089 | .030 | 159 | -2.987 | **.003** |
| Time | -.036 | .001 | 1919 | -43.381 | **<.001** |
| Infection*Time | .003 | .001 | 1919 | 2.319 | **.021** |
| **Three Doses vs. Zero Doses A (day 6 to day 30)** |  |  |  |  |  |
| **Fixed effect** | *Value* | *Std.Error* | *df* | *t-value* | *p-value* |
| (Intercept) | 1.091 | .027 | 1486 | 39.854 | **<.001** |
| Infection | .132 | .039 | 145 | 3.366 | **.001** |
| Time | -.038 | .001 | 1486 | -36.103 | **<.001** |
| Infection*Time | -.005 | .002 | 1486 | -3.303 | **.001** |
| **STIs vs. Zero Doses B (day 8 to day 30)** |  |  |  |  |  |
| **Fixed effect** | *Value* | *Std.Error* | *df* | *t-value* | *p-value* |
| (Intercept) | 1.066 | .029 | 1375 | 36.926 | **<.001** |
| Infection | .095 | .042 | 156 | 2.263 | **.025** |
| Time | -.036 | .001 | 1375 | -29.258 | **<.001** |
| Infection*Time | -.004 | .002 | 1375 | -2.361 | **.018** |
| **DTIs vs. Zero Doses B (day 6 to day 30)** |  |  |  |  |  |
| **Fixed effect** | *Value* | *Std.Error* | *df* | *t-value* | *p-value* |
| (Intercept) | 1.012 | .026 | 1293 | 39.605 | **<.001** |
| Infection | -.110 | .041 | 141 | -2.697 | **.008** |
| Time | -.034 | .001 | 1293 | -30.161 | **<.001** |
| Infection*Time | .002 | .002 | 1293 | .919 | **.358** |

**Table S2.** Results of the time-specific natural logarithm of mortality (*day 8 to day 80). Related to Figure 3E.

| **Three Doses vs. Zero Doses** |  |  |  |  |
| --- | --- | --- | --- | --- |
|  | *Estimate* | *Std.Error* | *t-value* | *p-value* |
| (Intercept) | -4.824 | .185 | -26.078 | **<.001** |
| Infection | .868 | .262 | 3.318 | **.002** |
| Time | .044 | .004 | 11.609 | **<.001** |
| Infection*Time | -.012 | .005 | -2.257 | **.031** |
| **STIs vs. Zero Doses** |  |  |  |  |
|  | *Estimate* | *Std.Error* | *t-value* | *p-value* |
| (Intercept) | -4.824 | .209 | -23.030 | **<.001** |
| Infection | .946 | .295 | 3.195 | **.003** |
| Time | .044 | .004 | 10.252 | **<.001** |
| Infection*Time | -.014 | .006 | -2.285 | **.029** |

\* The breakpoints were determined using Change-Point Detection Algorithms (see Muggeo 2017) and iterative testing in the R package 'Segmented', available at https://cran.rproject.org/web/packages/segmented/index.html.
In the comparison between Three dose and the Zero doses control, breakpoints at day 8.60 and day 79.85 exhibited convergent model and the highest R-squared value. Similarly, in the comparison between the STIs and the control, days 8.16 and 75.50 were estimated as breakpoints. Thus, the interval from day 8 to day 80 was selected for fitting the piecewise regression.
Muggeo, V. M. (2017). Interval estimation for the breakpoint in segmented regression: a smoothed score-based approach. *Australian & New Zealand Journal of Statistics*, 59(3), 311-322.

**Table S3.**
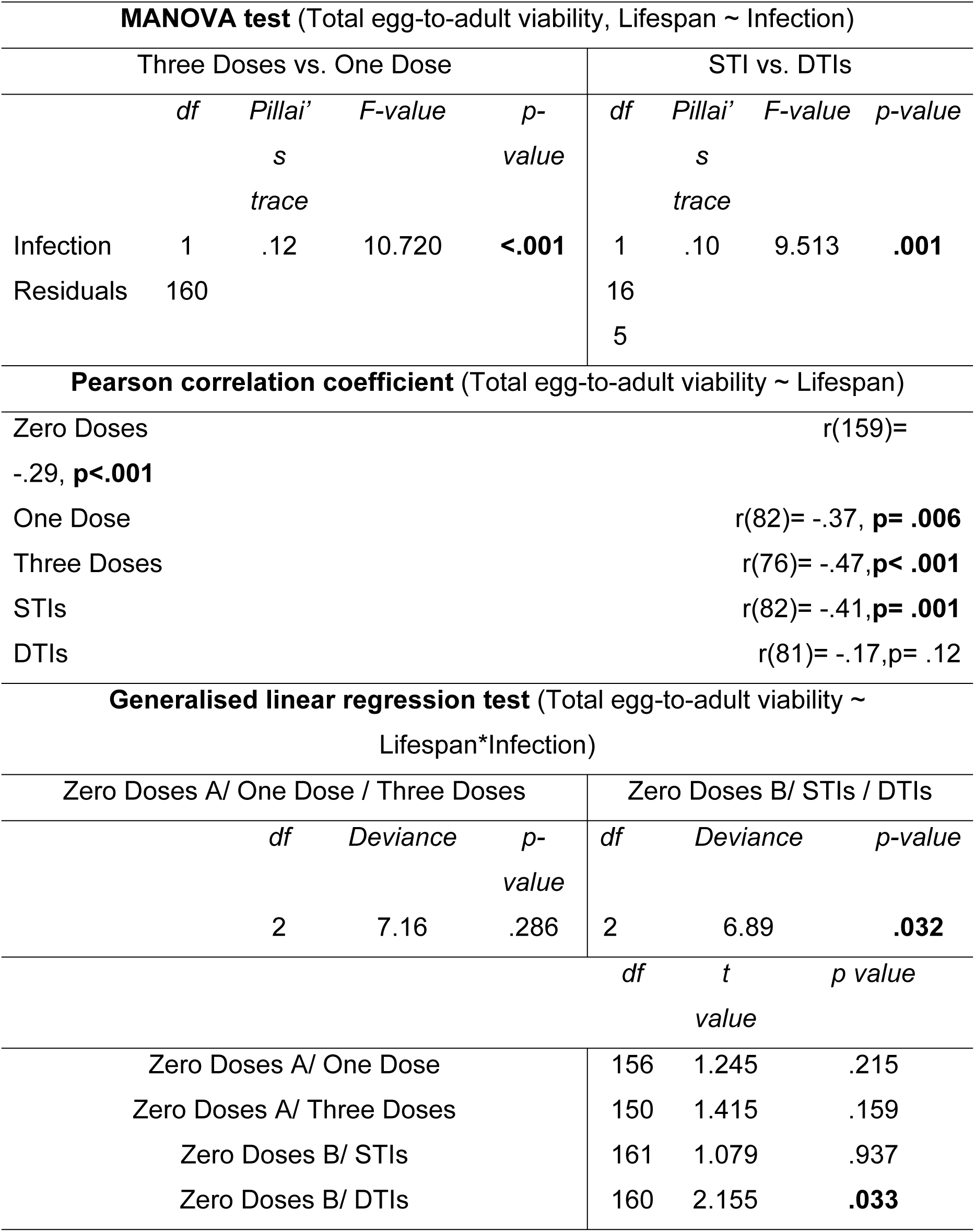
Results of effect of infection dose on determining life-history trade-offs between total viability and longevity. Related to Figure 4.

**Table S4.** Results of LME on age-specific reproduction among different genotypes following STIs. Related to Figure 5.

| **TotC expression** |  |  |  |  |  |
| --- | --- | --- | --- | --- | --- |
| **Fixed effect** | *Value* | *Std.Error* | *df* | *t-value* | *p-value* |
| (Intercept) | 52.129 | 4.534 | 932 | 11.497 | **<.001** |
| Infection | -3.256 | 6.537 | 230 | -.498 | .619 |
| Time | 7.617 | .789 | 932 | 9.651 | **<.001** |
| Genotype | -1.909 | 5.600 | 230 | -.341 | .734 |
| Infection*Time | .557 | 1.138 | 932 | .489 | .625 |
| Infection*Genotype | 13.177 | 8.031 | 230 | 1.641 | .102 |
| Time*Genotype | 2.260 | .975 | 932 | 2.319 | **.021** |
| Infection*Time*Genotype | -2.755 | 1.398 | 932 | -1.971 | **.049** |
| **Dif expression** |  |  |  |  |  |
| **Fixed effect** | *Value* | *Std.Error* | *df* | *t-value* | *p-value* |
| (Intercept) | 52.911 | 4.426 | 952 | 11.954 | **<.001** |
| Infection | -.390 | 6.414 | 235 | -.061 | .952 |
| Time | 8.330 | .740 | 952 | 11.252 | **<.001** |
| Genotype | -9.558 | 5.487 | 235 | -1.742 | .083 |
| Infection*Time | 1.004 | 1.073 | 952 | .936 | .349 |
| Infection*Genotype | 9.299 | 7.966 | 235 | 1.167 | .244 |
| Time*Genotype | .066 | .918 | 952 | .072 | .943 |
| Infection*Time*Genotype | -3.355 | 1.332 | 952 | -2.518 | **.012** |

**Table S5.** Results of differential contrast analysis of the effect of *TotC* and *Dif* expression on fecundity under STIs.

| *Estimate* | *Std. Error* | *df* | *t value* | *p-value\** |
| --- | --- | --- | --- | --- |
| -9.099 | 4.499 | 465 | -2.022 | **.044** |

\* The *p*-value was obtained through pairwise comparisons using the general linear hypothesis test.
And the model is:
where,
'kd' represents the knockdown genotype, 'ctrl' denotes the combined control genotype, and the acronyms 'STI', 'DTI', and 'NTR' stand for sexually transmitted infection, directly topical infection, and non-infection, respectively.

**Table S6.** Results of the differential contrast analysis of the effect of *Dif* expression on fecundity under STIs and DTIs. Related to Figure S3E.

| *Estimate* | *Std. Error* | *df* | *t value* | *p-value\** |
| --- | --- | --- | --- | --- |
| 182.29 | 30.51 | 486 | 5.974 | **<.001** |

\* The *p*-value was obtained through pairwise comparisons using the general linear hypothesis test.
And the model is:
where,
'kd' represents the knockdown genotype, 'ctrl' denotes the combined control genotype, and the acronyms 'STI', 'DTI', and 'NTR' stand for sexually transmitted infection, directly topical infection, and non-infection, respectively.

## References

1. Adamo, S. A. (2014), Parasitic Aphrodisiacs: Manipulation of the Hosts’ Behavioral Defenses by Sexually Transmitted Parasites, Integrative and Comparative Biology, 54(2), 159–165.

2. Agaisse, H., Petersen, U. M., Boutros, M., Mathey-Prevot, B., & Perrimon, N. (2003). Signaling role of hemocytes in Drosophila JAK/STAT-dependent response to septic injury. Developmental cell, 5(3), 441–450.

3. Amstrup, A. B., Bæk, I., Loeschcke, V., & Sørensen, J. G. (2022). A functional study of the role of Turandot genes in Drosophila melanogaster: An emerging candidate mechanism for inducible heat tolerance. Journal of Insect Physiology, 143, 104456.

4. An, S., Dong, S., Wang, Q., Li, S., Gilbert, L. I., Stanley, D., & Song, Q. (2012). Insect neuropeptide bursicon homodimers induce innate immune and stress genes during molting by activating the NF-κB transcription factor Relish. PLoS One, 7(3), e34510.

5. Bacalum, M., & Radu, M. (2015). Cationic antimicrobial peptides cytotoxicity on mammalian cells: an analysis using therapeutic index integrative concept. International Journal of Peptide Research and Therapeutics, 21, 47–55.

6. Bonneaud, C., Mazuc, J., Chastel, O., Westerdahl, H., & Sorci, G. (2004). Terminal investment induced by immune challenge and fitness traits associated with major histocompatibility complex in the house sparrow. Evolution, 58(12), 2823–2830.

7. Brannelly, L. A., Webb, R., Skerratt, L. F., & Berger, L. (2016). Amphibians with infectious disease increase their reproductive effort: evidence for the terminal investment hypothesis. Open Biology, 6(6), 150251.

8. Brommer, J. E. (2000). The evolution of fitness in life-history theory. Biological Reviews, 75(3), 377–404.

9. Clutton-Brock, T. H. (1984). Reproductive effort and terminal investment in iteroparous animals. The American Naturalist, 123(2), 212–229.

10. Corbel, Q., & Carazo, P. (2022). Perception of dead conspecifics increases reproductive investment in fruit flies. Functional Ecology, 36(8), 1834–1844.

11. Dong, S., & Dimopoulos, G. (2023). Aedes aegypti Argonaute 2 controls arbovirus infection and host mortality. Nature communications, 14(1), 5773.

12. Duffield, K. R., Bowers, E. K., Sakaluk, S. K., & Sadd, B. M. (2017). A dynamic threshold model for terminal investment. Behavioral Ecology and Sociobiology, 71, 1–17.

13. Ebert, D., & Mangin, K. L. (1997). The influence of host demography on the evolution of virulence of a microsporidian gut parasite. Evolution, 51(6), 1828–1837.

14. Ekengren, S., & Hultmark, D. (2001). A family of Turandot-related genes in the humoral stress response of Drosophila. Biochemical and Biophysical Research Communications, 284(4), 998–1003.

15. Farchmin, P. A., Eggert, A. K., Duffield, K. R., & Sakaluk, S. K. (2020). Dynamic terminal investment in male burying beetles. Animal Behaviour, 163, 1–7.

16. Foo, Y. Z., Lagisz, M., O’Dea, R. E., & Nakagawa, S. (2023). The influence of immune challenges on the mean and variance in reproductive investment: a meta-analysis of the terminal investment hypothesis. BMC Biology, 21(1), 107.

17. Hansen, T. F. (2003). Is modularity necessary for evolvability?: Remarks on the relationship between pleiotropy and evolvability. Biosystems, 69(2-3), 83–94.

18. Hanson, M. A., Grollmus, L., & Lemaitre B. (2023). Ecology-relevant bacteria drive the evolution of host antimicrobial peptides in Drosophila. Science 381(6655), eadg5725.

19. He, X., & Zhang, J. (2006). Toward a molecular understanding of pleiotropy. Genetics, 173(4), 1885–1891.

20. Hirshfield, M. F., & Tinkle, D. W. (1975). Natural selection and the evolution of reproductive effort. Proceedings of the National Academy of Sciences, 72(6), 2227–2231.

21. Hoffmann, J. A., & Reichart J. M. (2002). Drosophila innate immunity: an evolutionary perspective. Nature Immunology 3(2), 121–126.

22. Hong, S., Shang, J., Sun, Y., Tang, G., & Wang, C. (2023). Fungal infection of insects: molecular insights and prospects. Trends in Microbiology, 32(3), 302–316.

23. Houle, D., & Rossoni, D. M. (2022). Complexity, evolvability, and the process of adaptation. Annual Review of Ecology, Evolution, and Systematics, 53(1), 137–159.

24. Hudson, A. L., Moatt, J. P., & Vale, P. F. (2020). Terminal investment strategies following infection are dependent on diet. Journal of Evolutionary Biology, 33(3), 309–317.

25. Immonen, E., & Ritchie, M. G. (2012). The genomic response to courtship song stimulation in female Drosophila melanogaster. Proceedings of the Royal Society B: Biological Sciences, 279(1732), 1359–1365.

26. Ip, Y. T., Reach, M., Engstrom, Y., Kadalayil, L., Cai, H., González-Crespo, S., … & Levine, M. (1993). *Dif*, a dorsal-related gene that mediates an immune response in Drosophila. Cell, 75(4), 753–763.

27. Jehan, C., Sabarly, C., Rigaud, T., & Moret, Y. (2022). Age-specific fecundity under pathogenic threat in an insect: Terminal investment versus reproductive restraint. Journal of Animal Ecology, 91(1), 101–111.

28. Kirkwood, T. B., & Rose, M. R. (1991). Evolution of senescence: late survival sacrificed for reproduction. Philosophical Transactions of the Royal Society of London. Series B: Biological Sciences, 332(1262), 15–24.

29. Knell, R. J., Webberley, K. M. (2004) Sexually transmitted diseases of insects: distribution, evolution, ecology and host behaviour, Biological Reviews 79(3), 557–81.

30. Kolora, S. R. R., Owens, G. L., Vazquez, J. M., Stubbs, A., Chatla, K., Jainese, C., … & Sudmant, P. H. (2021). Origins and evolution of extreme life span in Pacific Ocean rockfishes. Science, 374(6569), 842–847.

31. Laskowski, K. L., Moiron, M., & Niemelä, P. T. (2021). Integrating behavior in life-history theory: allocation versus acquisition?. Trends in Ecology & Evolution, 36(2), 132–138.

32. Lemaitre, B., Nicolas, E., Michaut, L., Reichhart, J. M., & Hoffmann, J. A. (1996). The dorsoventral regulatory gene cassette spätzle/Toll/cactus controls the potent antifungal response in Drosophila adults. Cell, 86(6), 973–983.

33. Lemaître, J. F., Berger, V., Bonenfant, C., Douhard, M., Gamelon, M., Plard, F., & Gaillard, J. M. (2015). Early-late life trade-offs and the evolution of ageing in the wild. Proceedings of the Royal Society B: Biological Sciences, 282(1806), 20150209.

34. Lemaître, J. F., Moorad, J., Gaillard, J. M., Maklakov, A. A., & Nussey, D. H. (2024). A unified framework for evolutionary genetic and physiological theories of aging. PLoS Biology, 22(2), e3002513.

35. Liao, A., Cavigliasso, F., Savary, L., & Kawecki, T. J. (2024). Effects of an entomopathogenic fungus on the reproductive potential of Drosophila males. Ecology and Evolution, 14(4), e11242.

36. Lu, H. L., & St. Leger, R. J. (2016). Insect immunity to entomopathogenic fungi. Advances in Genetics, 94, 251–285.

37. Martins, N. E., Faria, V. G., Teixeira, L., Magalhaes, S., & Sucena, E. (2013). Host adaptation is contingent upon the infection route taken by pathogens. PLoS Pathogens, 9(9): e1003601.

38. McKean, K. A., Yourth, C. P., Lazzaro, B. P., & Clark, A. G. (2008). The evolutionary costs of immunological maintenance and deployment. BMC evolutionary biology, 8, 1–19.

39. McKean, K. A., & Lazzaro, B. P. (2011). The costs of immunity and the evolution of immunological defense mechanisms. Mechanisms of life history evolution, 299–310.

40. Minchella, D. J., & Loverde, P. T. (1981). A cost of increased early reproductive effort in the snail Biomphalaria glabrata. The American Naturalist, 118(6), 876–881.

41. Moita, L. F., Wang-Sattler, R., Michel, K., Zimmermann, T., Blandin, S., Levashina, E. A., & Kafatos, F. C. (2005). In vivo identification of novel regulators and conserved pathways of phagocytosis in *A. gambiae*. Immunity, 23(1), 65–73.

42. Ortiz-Urquiza, A., & Keyhani, N. O. (2013). Action on the surface: entomopathogenic fungi versus the insect cuticle. Insects, 4(3), 357–374.

43. 43. Parker, B. J., Barribeau, S. M., Laughton, A. M., de Roode, J. C., & Gerardo, N. M. (2011). Non-immunological defense in an evolutionary framework. Trends in Ecology & Evolution, 26(5), 242–248.

44. Pianka, E. R., & Parker, W. S. (1975). Age-specific reproductive tactics. The American Naturalist, 109(968), 453–464.

45. Pinheiro, J., Bates, D., DebRoy, S., and Sarkar, D. (2012). Nonlinear mixed-effects models. R package version, 3, 1–89.

46. Polak, M., & Starmer, W. T. (1998). Parasite-induced risk of mortality elevates reproductive effort in male Drosophila. Proceedings of the Royal Society B: Biological Sciences. 265(1411), 2197–2201.

47. R Core Team, 2014. R: A language and environment for statistical computing. R Foundation for Statistical Computing, Vienna, Austria.

48. Rafaluk-Mohr, C., Gerth, M., Sealey, J. E., Ekroth, A. K., Aboobaker, A. A., Kloock, A., & King, K. C. (2022). Microbial protection favors parasite tolerance and alters host-parasite coevolutionary dynamics. Current Biology, 32(7), 1593–1598.

49. Rauw, W. M. (2012). Immune response from a resource allocation perspective. Frontiers in genetics, 3, 267.

50. Reznick, D., Nunney, L. and Tessier, A., 2000. Big houses, big cars, superfleas and the costs of reproduction. Trends in ecology & evolution, 15(10), pp.421–425.

51. Roberts, D. W., & St Leger, R. J. (2004). Metarhizium spp., cosmopolitan insect-pathogenic fungi: mycological aspects. Advances in applied microbiology, 54(1), 1–70.

52. Rommelaere, S., Carboni, A., Juarez, J. F. B., Boquete, J. P., Abriata, L. A., Meireles, Teixeira Pinto Meireles, F., Rukes, V., Vincent, C., Kondo, S., Dionne, M. S., Dal Peraro, M., Cao, C., & Lemaitre, B. (2024). A humoral stress response protects Drosophila tissues from antimicrobial peptides. Current Biology, 34(7), 1426–1437.

53. Saraux, C., & Chiaradia, A. (2022). Age-related breeding success in little penguins: a result of selection and ontogenetic changes in foraging and phenology. Ecological monographs, 92(1), e01495.

54. Schneider, D. S., & Ayres, J. S. (2008). Two ways to survive infection: what resistance and tolerance can teach us about treating infectious diseases. Nature Reviews Immunology, 8(11), 889–895.

55. Schulz, N. K., Stewart, C. M., & Tate, A. T. (2023). Female investment in terminal reproduction or somatic maintenance depends on infection dose. Ecological Entomology, 48(6), 714–724.

56. Schwenke, R. A., Lazzaro, B. P., & Wolfner, M. F. (2016). Reproduction– immunity trade-offs in insects. Annual review of entomology, 61(1), 239–256.

57. Stearns, S. C. (1992). The Evolution of Life Histories. Oxford University Press.

58. Stearns, F. W. (2010). One hundred years of pleiotropy: a retrospective. Genetics, 186(3), 767–773.

59. Than, A. T., Ponton, F., & Morimoto, J. (2020). Integrative developmental ecology: a review of density-dependent effects on life-history traits and host-microbe interactions in non-social holometabolous insects. Evolutionary Ecology, 34(5), 659–680.

60. Thomas, F., Adamo, S. S., & Moore, J. (2005). Parasitic manipulation: where are we and where should we go?. Behavioural processes, 68(3), 185–199.

61. Van Noordwijk, A. J. and De Jong, G., 1986. Acquisition and allocation of resources: their influence on variation in life history tactics. The American Naturalist, 128(1), pp.137–142.

62. 62. Van Rheenen, W., Peyrot, W. J., Schork, A. J., Lee, S. H., & Wray, N. R. (2019). Genetic correlations of polygenic disease traits: from theory to practice. Nature Reviews Genetics, 20(10), 567–581.

63. Viney, M. E., Riley, E. M., & Buchanan, K. L. (2005). Optimal immune responses: immunocompetence revisited. Trends in ecology & evolution, 20(12), 665–669.

64. Weil, Z. M., Martin, L. B., Workman, J. L., & Nelson, R. J. (2006). Immune challenge retards seasonal reproductive regression in rodents: evidence for terminal investment. Biology Letters, 2(3), 393–396.

65. Wigby, S., Domanitskaya, E. V., Choffat, Y., Kubli, E., & Chapman, T. (2008). The effect of mating on immunity can be masked by experimental piercing in female Drosophila melanogaster, Journal of Insect Physiology, 54(2), 414–420.

66. Williams, G. C. (1957). Pleiotropy, Natural Selection, and the Evolution of Senescence. Evolution, 11(4), 398–411.

67. Williams, G. C. (1966). Natural selection, the costs of reproduction, and a refinement of Lack’s principle. The American Naturalist, 100(916), 687–690.

68. Zerofsky, M., Harel, E., Silverman, N., & Tatar, M. (2005). Aging of the innate immune response in Drosophila melanogaster. Aging cell, 4(2), 103–108.

69. Zhong, W., McClure, C. D., Evans, C. R., Mlynski, D. T., Immonen, E., Ritchie, M. G., & Priest, N. K. (2013). Immune anticipation of mating in Drosophila: Turandot M promotes immunity against sexually transmitted fungal infections. Proceedings of the Royal Society B: Biological Sciences, 280(1773), 20132018.

